# New topologies in the unfolding of the Doubly Degenerate Bogdanov-Takens bifurcation

**DOI:** 10.1101/2025.11.02.686129

**Authors:** Marisa Saggio

## Abstract

High-codimension bifurcations play a key role in shaping the dynamics of nonlinear models, as their unfoldings establish structured relationships between lower-codimension bifurcations and, ultimately, the attractors they generate. Owing to this unifying and predictive capacity, such bifurcations are attracting growing interest in mathematical biology, particularly in neuroscience. One notable example is the codimension-3 Degenerate Bogdanov-Takens (DBT) bifurcation, proposed as an organizing center for neural dynamics and playing a key role in bursting. The DBT itself arises within the unfolding of an even higher-order bifurcation, the Doubly Degenerate Bogdanov-Takens (DDBT), whose structure remains only partially understood, despite initial indications of its relevance in neural dynamics. In this work, we expand the numerical investigation of the DDBT unfolding through the use of spherical surfaces, corroborating the conjecture that it connects DBT to a highly symmetric codimension-3 bifurcation, but through transitions that partly differ from those proposed in the literature. Notably, one key intermediate passage remains unsolved. We then show that using planes to explore the unfolding allows for new bifurcation topologies and transitions that cannot be found on spheres. We illustrate how transitions to topologies supporting excitability and fold/homoclinic bursting, two biologically relevant behaviors, differ across spheres and planes. Overall, the additional bifurcation structures enrich our understanding of what models with a DDBT bifurcation can do and how these dynamics are related.

## 1 Introduction

When describing a system using differential equations, we often focus on the stable configurations it eventually settles into: the *attractors* of the dynamics. As system parameters change, these attractors can appear, disappear, or switch stability. This qualitative transformation of the attractor landscape is known as a *bifurcation*, and a bifurcation diagram depicts these changes under parameter variation. Parameters may represent, for example, internal properties or external conditions. Common bifurcations in planar systems (Fig. 1A), such as Fold (F), Hopf (H), Homoclinic (Hom), Fold of Limit Cycles (FLC) or Saddle-Node-on-Invariant-Circle (SNIC), require the variation of one parameter, and are thus called *codimension-1* (codim 1) bifurcations. If the system involves more parameters, we can study how codim 1 bifurcations depend on them, creating higher-dimensional bifurcation diagrams. For example, in a two-parameter bifurcation diagram, codim 1 bifurcations are curves, which stratify the space into regions within which the repertoire of available behaviors stays *qualitatively* the same (Fig. 1B). Bifurcation diagrams, thus, provide condensed information about what a system can do and how this varies with parameters [42].

**Fig. 1.**
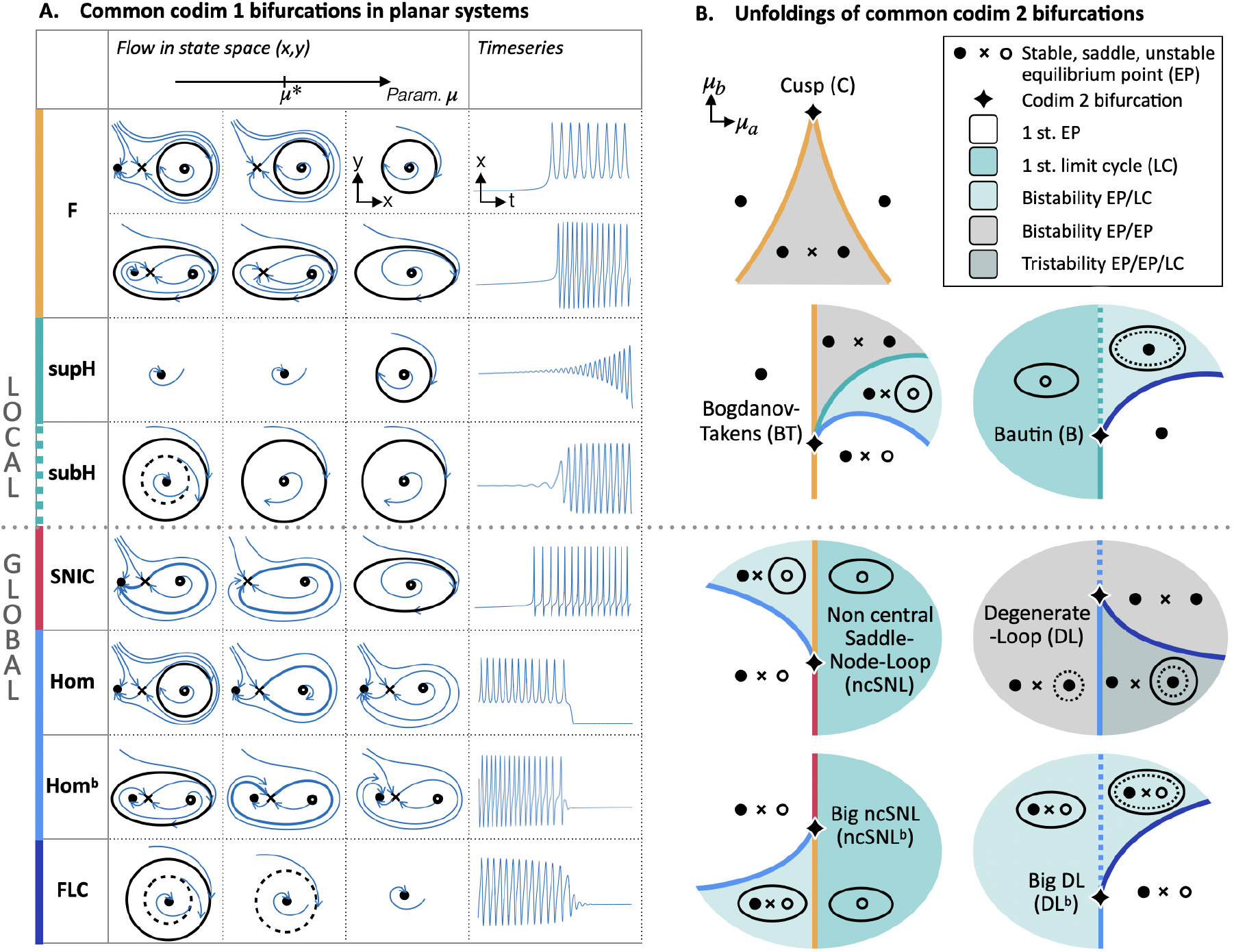
Common codim 1 and codim 2 bifurcations in planar dynamical systems. See, for example, [42]. **A**. Codim 1 bifurcations (abbreviations in Table 1) involving equilibria and limit cycles (stable/unstable sketched as full/dashed ellipses). For each bifurcation, we show the configuration of attractors and repellers in state space before, at (*µ*^*\**^), and after the bifurcation, and an example of a timeseries. The direction of the parameter change is arbitrary. Orbits in state space are shown in blue. Thicker curves identify homoclinic orbits (returning to the same equilibrium) and heteroclinic ones (connecting two equilibria). At a fold bifurcation, two equilibria coalesce and disappear, and the system can settle into another attractor (two examples are shown with the other attractor being a limit cycle). At a Hopf, the equilibrium stability changes and a limit cycle appears. At a SNIC, the fold is accompanied by the birth of a limit cycle. At a homoclinic, a limit cycle meets a saddle and disappears (also in the Hom^*b*^ variation, where the superscript *b* indicates a *big* limit cycle). At a FLC, two limit cycles coalesce and disappear. In supercritical/subcritical Hopf, SNIC and Hom bifurcations, the new object created is stable/unstable (only shown for the Hopf). **B**. Unfoldings of common codim 2 bifurcations. Codim 1 bifurcation curves stratify the parameter space (*µ*_*a*_, *µ*_*b*_). In each region, a possible state space configuration is shown. BT and ncSNL bifurcations can also involve subcritical homoclinic, Hopf and SNIC (not shown). NcSNL and DL are also presented in the variation with a big limit cycle. Bifurcations of equilibria (even though they can involve limit cycles) are often referred to as *local* bifurcations (top panels), while *global* bifurcations cannot be identified by looking at a small neighborhood of the equilibrium [42]).

Codim 1 bifurcations in two-parameter diagrams cannot occur in arbitrary configurations. Instead, they are arranged in characteristic ways, locally organized by *codim 2* bifurcations, that is, bifurcations requiring the precise tuning of two parameters. At a codim 2 bifurcation, two or more bifurcations coincide. Moving away in parameter space from the codim 2 bifurcation point, codim 1 curves unfold from it stereotypically, as described by the *unfolding* of that specific codim 2 bifurcation (Fig. 1B).

The configurations of codim 2 bifurcations themselves can be organized by the unfoldings of codim bifurcations, and so on. This implies that high-codimension bifurcations, far from being esoteric mathematical objects [41], act as organizing centers that, through their unfoldings, unify a system’s behaviors [28]. High-codimension bifurcations are the reason for which “there are some two-dimensional bifurcation diagrams of planar vector fields that seem to have a *universal character* in the sense that they occur in many models” [39]. Knowing that a system possesses a high-codimension bifurcation is therefore a compact yet powerful source of information about its possible dynamics, from lower-order bifurcations down to attractors. Recognizing in a two-parameter bifurcation diagram the signatures of a high-codimension bifurcation, on the other hand, can immediately reveal other bifurcation topologies that can be expected if parameters are varied, solely through the knowledge of the unfolding.

While many codim 1 and 2 bifurcations and their unfoldings are described in textbooks [42], this is rarely the case for higher-order bifurcations. Among them, the Degenerate Bogdanov-Takens (DBT) [20, 43], is a well-studied codim 3 bifurcation existing in four types and occurring in applications including ecology [5, 42], chemistry [10, 14] and neural dynamics. In neuroscience, it is present in several single neuron models [29, 1], and, in particular, in any conductance-based model for Type I excitability [40, 47, 62, 22, 23, 55, 16, 52, 25]. One type of DBT, the DBT-*focus*, has been proposed as an organizing center for the activity of single neurons [50, 40] and, possibly, for neural population dynamics [59, 56]. When coupled with slow variables to form a *fast–slow system*, it can produce several classes of bursting, some of which are observed in excitable cells [7, 26, 49, 56].

This codim 3 bifurcation is itself part of the unfolding of the codim 4 Doubly Degenerate Bogdanov-Takens (DDBT) bifurcation, whose structure remains not completely understood [39]. Notably, Osinga and colleagues showed that the known portion of this unfolding supports a type of bursting, fold/subH [60] (bursting with a fold bifurcation at the onset of oscillations and a subH at their offset), also observed in some excitable cells, which could not be explained by the DBT unfolding [49]. This led the authors to propose the DDBT, rather than simply the DBT, as an organizing center for excitable cells’ dynamics. The systematic analysis of which fast-slow bursters can be obtained by slowly moving in this unfolding (*unfolding-theory approach to fast–slow bursting* [26]) later identified paths for all sixteen theoretically possible planar point-cycle bursters [56, 34].

**Table 1.** List of abbreviations. Note that, while SNIC and SNL are synonyms, we use the first for the codim 1 bifurcation and the second for the non-central codim 2 one, as is common in the neuroscience literature.

| Abbr. | Meaning and some alternative names |
| --- | --- |
| **Solutions** |  |
| EP | Equilibrium Point (Fixed Point [58]) |
| LC | Limit Cycle |
| NS | Neutral-Saddle |
| **Codimension-1** |  |
| F | Fold (Saddle-Node, Limit-Point) |
| SNIC | Saddle-Node on Invariant Circle (Saddle-Node Loop) |
| Hom | Homoclinic (Saddle-Homoclinic, Saddle Loop, Saddle-Connection, Saddle Separatrix Loop) |
| H | Hopf (Andronov-Hopf) |
| FLC | Fold of Limit Cycles (Limit Point of Cycles, Double Limit Cycle) |
| **Codimension-2** |  |
| C | Cusp [42]<br>(codim 2 semi-hyperbolic node [39]) |
| BT | Bogdanov-Takens<br>(double-zero bifurcation [42], nilpotent cusp of order two) |
| B | Bautin (gen. Hopf [42], deg. Hopf-Takens [20]) |
| ncSNL | non-central Saddle-Node Loop<br>(Saddle-Node-Loop [13, 31], codim 2 Saddle-Node-Loop [20], codim 2 Saddle-Node-Homoclinic orbit [35], codim 2 Saddle-Node separatrix loop [33, 34]) |
| DL | Deg. Loop [20] (Separatrix loop with zero saddle value [57], a type of Resonant Homoclinic [11, 32]) |
| **Codimension-3 and 4** |  |
| DBT | Deg. Bogdanov-Takens - four types (focus, elliptic, saddle, cusp) [19, 20] |
| DBT-focus | Deg. Bogdanov-Takens focus type (Codim 3 nilpotent focus [39], Bogdanov-Takens-Cusp [40], cubic BT, nilpotent cusp of order three [41]) |
| DBT-cusp | Deg. Bogdanov-Takens cusp type (nilpotent cusp of order three [39], DBT with double equilibrium [43] cusp of codim 3 [43]) |
| CLC | Cusp of Limit Cycles |
| DDBT | Doubly Deg. Bogdanov-Takens |
| **Other** |  |
| sub/sup | subcritical/supercritical |
| gen. | generalized |
| deg. | degenerate |

In addition to the three unfolding parameters of the DBT bifurcation, the DDBT unfolding also considers the variation of a fourth parameter, called *b*, which is instead fixed and non-zero in the DBT case. The case in which *b* = 0 has been studied in detail [39] and gives a reflectionally symmetric bifurcation structure, here referred to as the *symmetric case*, found in models of electrical circuits, in chemistry, biochemistry and ecology [4, 38, 61, 8, 6, 45, 39]. In 1998, Khibnik, Krauskopf and Rousseau conjectured a number of intermediate bifurcation diagrams which, as *b* is increased from 0, modify the symmetric case and lead to the DBT-focus one for *b* sufficiently big [39]. This sequence is a representation of the unfolding of the codim 4 DDBT. In 2016, Krauskopf and Osinga investigated the cases closer to the DBT-focus case, which differed from those previously conjectured [41].

Given the interest in this bifurcation and its potential role in neural dynamics [40, 1, 48, 49], but also in other contexts such as cosmology [2] and fusion plasma transport models [15], we here expand the numerical study of its unfolding and provide some additional topologies that can be found when the unfolding is sliced with two-dimensional surfaces.

In the remainder of the Introduction, we briefly review the standard approach of using spherical surfaces to explore the unfolding, and describe the DBT-focus [20] and the symmetric cases [39]. A more detailed mathematical treatment can be found in [20, 39, 49, 41, 43]. Figures 1 and 5 serve as a quick reference for the well-known codim 1 and 2 bifurcations involved in this unfolding. In the Results section, we present our numerical explorations using spheres and discuss how they relate to the conjecture, highlighting differences and still unresolved transitions. We then show that using planes in parameter space, rather than spherical sections, reveals new bifurcation topologies potentially relevant for models in neuroscience and other systems where the DDBT has been identified. We discuss, in particular, how spheres and planes identify different transitions to excitable behaviors close to a SNIC, a distinguishing feature of neuronal Type I excitability [54, 35, 51], and fold/hom bursting.

### 1.1 Equations of the unfolding of the DDBT

The equations for the candidate unfolding of the DDBT are^1^ [39, 20]:

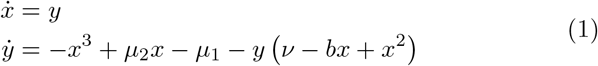

where the dot indicates *d/dt* and (*µ*_1_, *µ*_2_, *v, b*) are the unfolding parameters.

Equilibrium points, satisfying 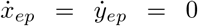, do not depend on *b*. Local bifurcations can be identified by imposing their specific defining conditions (as many as their codimension) and verifying that the non-degeneracy and transversality conditions hold. Defining/non-degeneracy conditions are expressed as the vanishing/non-vanishing of specific combinations of partial derivatives of the vector field at the equilibrium.

If we consider the Jacobian matrix of the system computed at its equilibria:

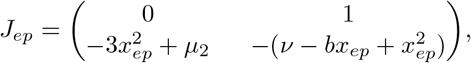

the manifold of fold bifurcation, at which the Jacobian has a single zero eigenvalue (defining condition), can be found by imposing det(*J*_*ep*_) = *λ*_1_*λ*_2_ = 0, where *λ*_1,2_ are the eigenvalues of *J*_*ep*_. In our case, it requires 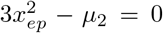, making the fold bifurcation manifold independent of *b*.

The two surfaces of fold bifurcation meet at (*µ*_1_, *µ*_2_) = (0, 0), where (*x*_*ep*_, *y*_*ep*_) = (0, 0). This is the codim 2 cusp (C) bifurcation where the three branches of equilibria coincide (Fig. 2A, B top), and corresponds to a higher order degeneracy of the equilibrium.

**Fig. 2.**
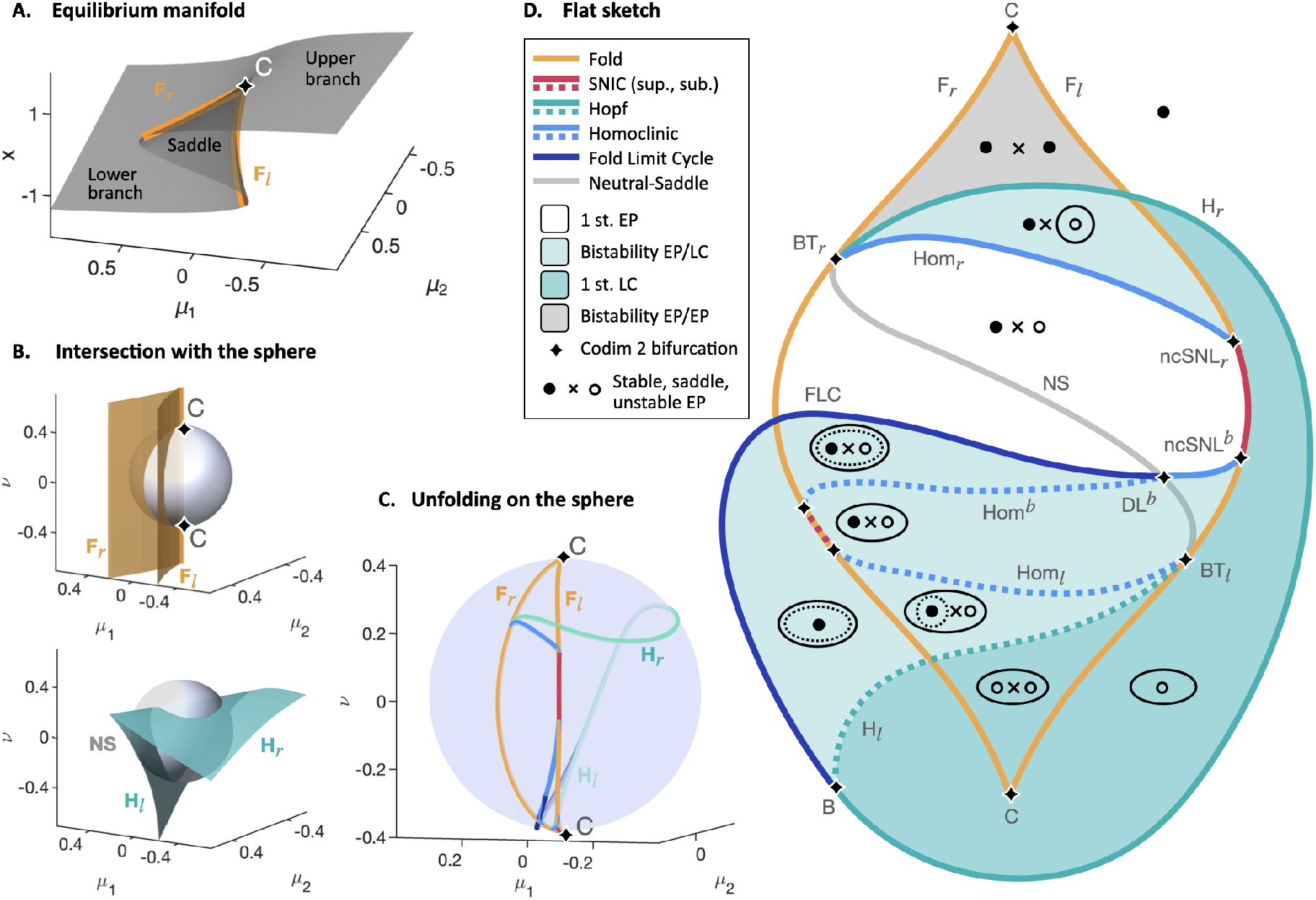
Unfolding of the degenerate Bogdanov-Takens focus on a spherical surface (b = 1). Summary of results in [20]. **A**. Equilibrium manifold, which depends only on *µ*_1_ and *µ*_2_, with curves of codim 1 fold bifurcations meeting at a codim 2 cusp point. **B**. In the (*µ*_1_, *µ*_2_, *v*) parameter space, there are surfaces of codim 1 bifurcations and curves of codim 2 bifurcations. Intersection between the spherical surface and the fold (top) or Hopf (bottom) surfaces are curves of codim 1 bifurcations; intersection with the cusp line are points of codim 2 bifurcations. **C**. Bifurcation diagram on the semi-transparent sphere, including the curves of intersection with the other codim 1 bifurcation surfaces. **D**. Flat topologically equivalent sketch of the bifurcation diagram. A colored background marks the type of attractors in each region. The codim 1 curves are: the fold F_*r*_/F_*l*_ creating the two rightmost/leftmost equilibria and meeting at two codim 2 C; the Hopf (H), joining the two codim 2 BT_*r*_, BT_*l*_, creating or destroying a limit cycle (for convenience, we separated it into two branches H_*r*_ and H_*l*_). Homoclinic curves Hom_*r*_/Hom_*l*_ are also stemming from the BT points, creating or destroying a limit cycle around the rightmost/leftmost equilibrium; Hom^*b*^ is responsible for a big limit cycle enclosing all three equilibria; FLC connects a codim 2 Bautin point on the H curve with a codim 2 DL^*b*^; SNIC curves join either the codim 2 ncSNL_*r*_ and ncSNL^*b*^ points or the analogous points on F_*r*_ (not labeled).

At a Hopf bifurcation, the Jacobian has a pair of purely imaginary eigenvalues. It requires tr(*J*_*ep*_) = *λ*_1_ + *λ*_2_ = 0, implying 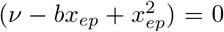, which depends on *b*. It also requires det(*J*_*ep*_) *>* 0. When det(*J*_*ep*_) *<* 0, the equilibria are saddles, and tr(*J*_*ep*_) = 0 identifies the Neutral-Saddle (NS) manifold. Neutral-saddle is not a bifurcation, but helps locate codim 2 DL points.

When both det(*J*_*ep*_) = tr(*J*_*ep*_) = 0, but *J*_*ep*_ *≠* 0, we have the codim 2 Bogdanov-Takens (BT) bifurcation, a double-zero eigenvalue bifurcation, dependent on *b* because of the condition on the trace.

A codim 2 BT bifurcation needs to satisfy two additional non-degeneracy conditions. Their violation leads to either a codim 3 DBT bifurcation with a triple equilibrium, which exists in saddle, focus and elliptic types [20], or with a double equilibrium, called the cusp type [19]. The DBT-focus type occurs at (*µ*_1_, *µ*_2_, *v*) =(0, 0, 0) and 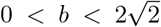, and the elliptic type for 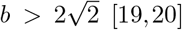. The saddle case requires a positive coefficient for *x*^3^ in Eq. 1.

The choice of signs in Eq. 1 is such that it has a stable DBT-focus [20]. This supports the greatest amount of fast/slow bursting patterns [56]. In addition, Eq. 1 gives a manifold of equilibria (Fig. 2A) in which the lower branch is mostly stable, while oscillations can surround the upper branch. Such a configuration is suitable, for example, to build bursting systems in which the active oscillatory phase is at a higher value of *x* as compared to the rest solution, as is typical in excitable cells. The reverse, as can be observed, for example, during seizure-like events [37], can be achieved by applying the transformation *x → −x* to Eq. 1. Bifurcation diagrams as those shown in this paper stay unchanged under this scenario, while state space representation for each region needs to be flipped horizontally.

Finally, the violation of both of these BT’s non-degeneracy conditions gives the codim 4 DDBT [39, 13, 49]. This occurs at the origin of the state space and for all parameters being equal to zero.

Relationships between the local bifurcations involved in this work are summarized in Fig. 3.

**Fig. 3.**
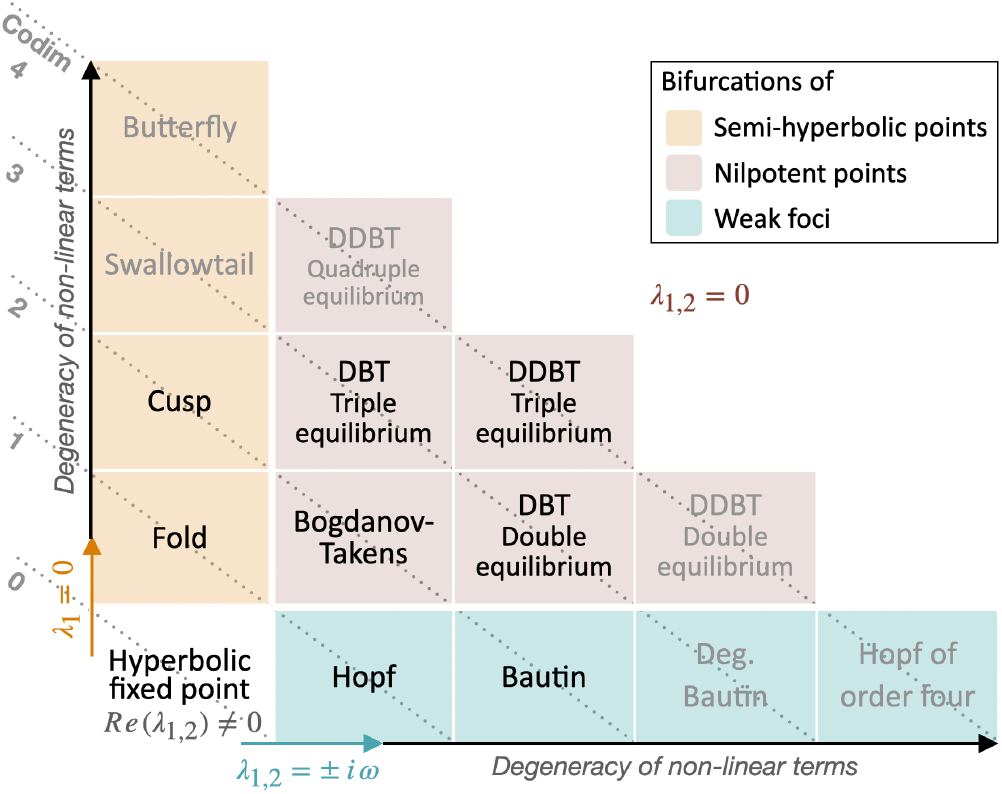
Review of the degeneracies of equilibria involved in local bifurcations in planar systems. A hyperbolic equilibrium point is one for which the eigenvalues of the Jacobian have nonzero real parts. Its stability is robust under small perturbations. In contrast, a non-hyperbolic equilibrium (*singularity*) is structurally unstable and can undergo bifurcations when parameters are varied [58, 42]. Non-hyperbolicity occurs when a single eigenvalue is zero (semihyperbolic equilibrium) [46]; when the eigenvalues are purely imaginary (weak focus) [27]; or when both eigenvalues are zero (nilpotent point, with non-trivial *J*_*ep*_ ≠ 0) [18, 39, 42]. The degree of degeneracy of the non-linear terms within each family leads to bifurcations of increasing codimension. In the progression, non-degeneracy conditions for a bifurcation are violated and this becomes part of the defining conditions for the next. BT can be conceptualized as a bifurcation point where fold and Hopf bifurcations co-occur [49]. The degenerate BT (DBT) exists in four types, three with triple equilibrium, and one with double equilibrium, depending on which BT non-degeneracy conditions are violated. They can be thought of as a BT bifurcation occurring on a cusp point [49], or on a Bautin (i.e., degenerate Hopf) point, respectively. The doubly degenerate BT (DDBT) singularity also exists in different types; we here work with the triple equilibrium focus type [20, 39, 18]. Its unfolding contains the bifurcations in black. Other local bifurcations fitting this table are written in gray. Codim 4 nilpotent bifurcations are also possible with double equilibrium (the cusp of order 4) [44, 64] and quadruple equilibrium (the less understood nilpotent saddle-node singularity of order 4) [18, 65]. The unfoldings of the bifurcations related to local singularities can contain global bifurcations.

### 1.2 Using spheres to explore the unfolding

#### Case 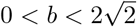: the unfolding of the DBT-focus

As explained above, for any positive 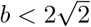, a codim 3 DBT-focus is found when the other three parameters are all equal to zero. Moving away from the origin of the (*µ*_1_, *µ*_2_, *v*) space, two-dimensional manifolds of codim 1 bifurcations emerge, with curves of codim 2 bifurcations at some of their intersections. For example, In Fig. 2B, the coincidence of H_*r*_ and F_*r*_ at BT_*r*_ is a codim 2 bifurcation because the two bifurcation conditions are verified at the same equilibrium, whereas that between H_*r*_ and F_*l*_ is not, as the two conditions involve different equilibria (we will adopt a softer definition when discussing codim 3 events).

It is not easy to deal with a 3-dimensional space. We can thus consider the intersection between the surface of a sphere centered at the origin (that is, on the DBT-focus) and the manifolds of codim 1 and codim 2 bifur-cations (Fig. 2B,C,D). This provides a two-dimensional bifurcation diagram for each radius *R* = const, where the angles of the spherical coordinates (*θ, ϕ*) are the parameters. How useful is this reduced diagram for capturing the unfolding properties? It can be shown [20, 39], that for sufficiently small values of *R*, the topology of the bifurcation diagram on the sphere stays the same, therefore providing an informative representation of the unfolding of the DBT-focus. More precisely, for any *b*^*\**^ *>* 0, there exists a sufficiently small value of the radius, *R*^*\**^, such that the same bifurcation topology can be found on any sphere with *R < R*^*\**^, while the topol-ogy changes on bigger spheres (Fig. 4A). A bifurcation diagram as such is said to have a *cone structure* in a neighborhood of the origin [41].

**Fig. 4.**
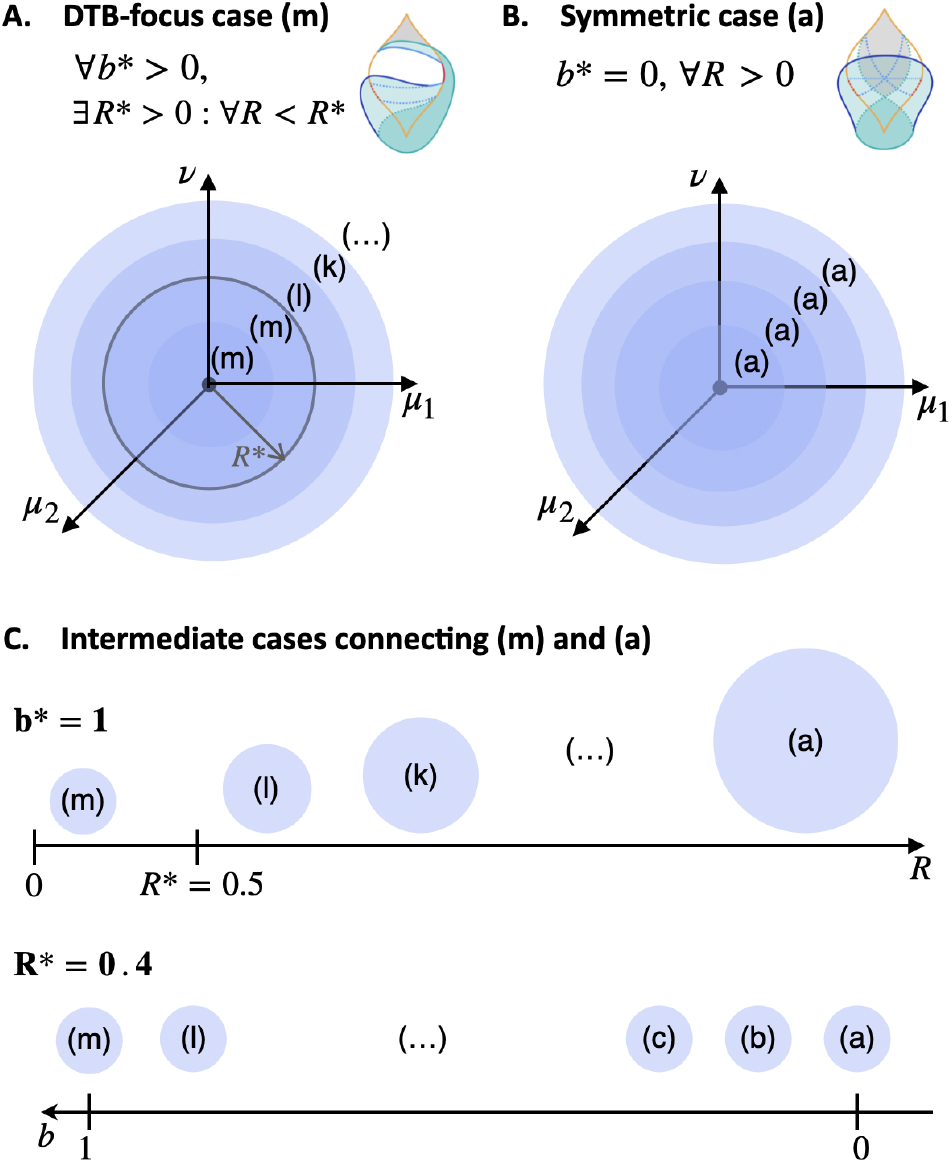
Exploring the four-dimensional unfolding using spheres with either fixed radius *R* or fixed *b*. Concepts described in [41]. **A**. The bifurcation diagram of the DBT-focus case (m) can be found on sufficiently small spheres for any fixed positive value of *b*. For bigger spheres, the topology of the bifurcation diagram goes through a series of different cases (l,k, …). **B**. When *b* = 0, the symmetric case (a) appears on any sphere. **C**. We can study the sequence of cases connecting (a) to (m) by fixing *b* and increasing *R* (top), or vice versa (bottom, the approach taken here). Values in the figure are those used in this paper.

The bifurcation diagram of the unfolding of the DBT-focus (Fig. 2C,D) is very rich. The bifurcations it includes are reviewed in Figures 1 and 5.

**Fig. 5.**
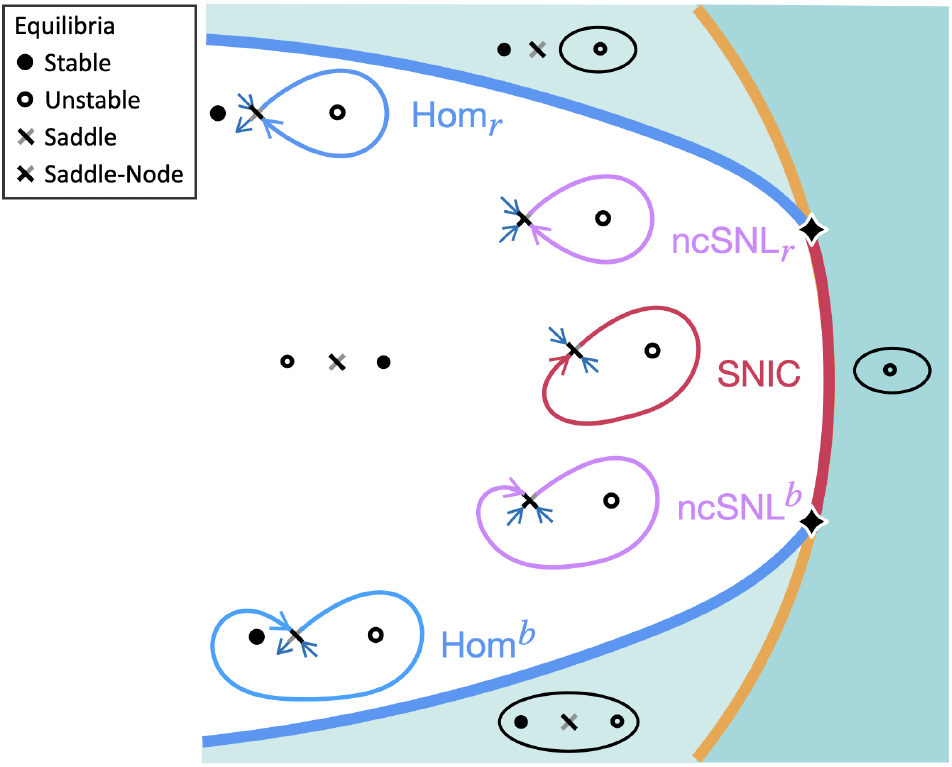
Summary of some bifurcations of homoclinic orbits in the unfolding. State space diagrams drawn also at the bifurcation curves/points [20, 42], with homoclinic orbits to the saddle or saddle-node shown as thick colored curves. Homoclinic orbits can return to the equilibrium point along different directions, shown in black in the saddle/saddle-node markers. For the saddle, the two possible return directions lie on its stable manifold and are related to the codim-1 bifurcation curves Hom and Hom^*b*^. For the saddle-node, the homoclinic orbit can return along the saddle-node center manifold, associated with the codim 1 SNIC bifurcation curve, or along non-central directions, giving the codim 2 ncSNL and ncSNL^*b*^ bifurcation points.

#### Case b = 0: unfolding the symmetric case

When *b* = 0, the bifurcation diagram has cone structure in the full space [39]: bifurcation diagrams on spheres are topologically equivalent for any *R* (Fig. 4B). This bifurcation diagram (Fig. 6) is the main focus of [39].

**Fig. 6.**
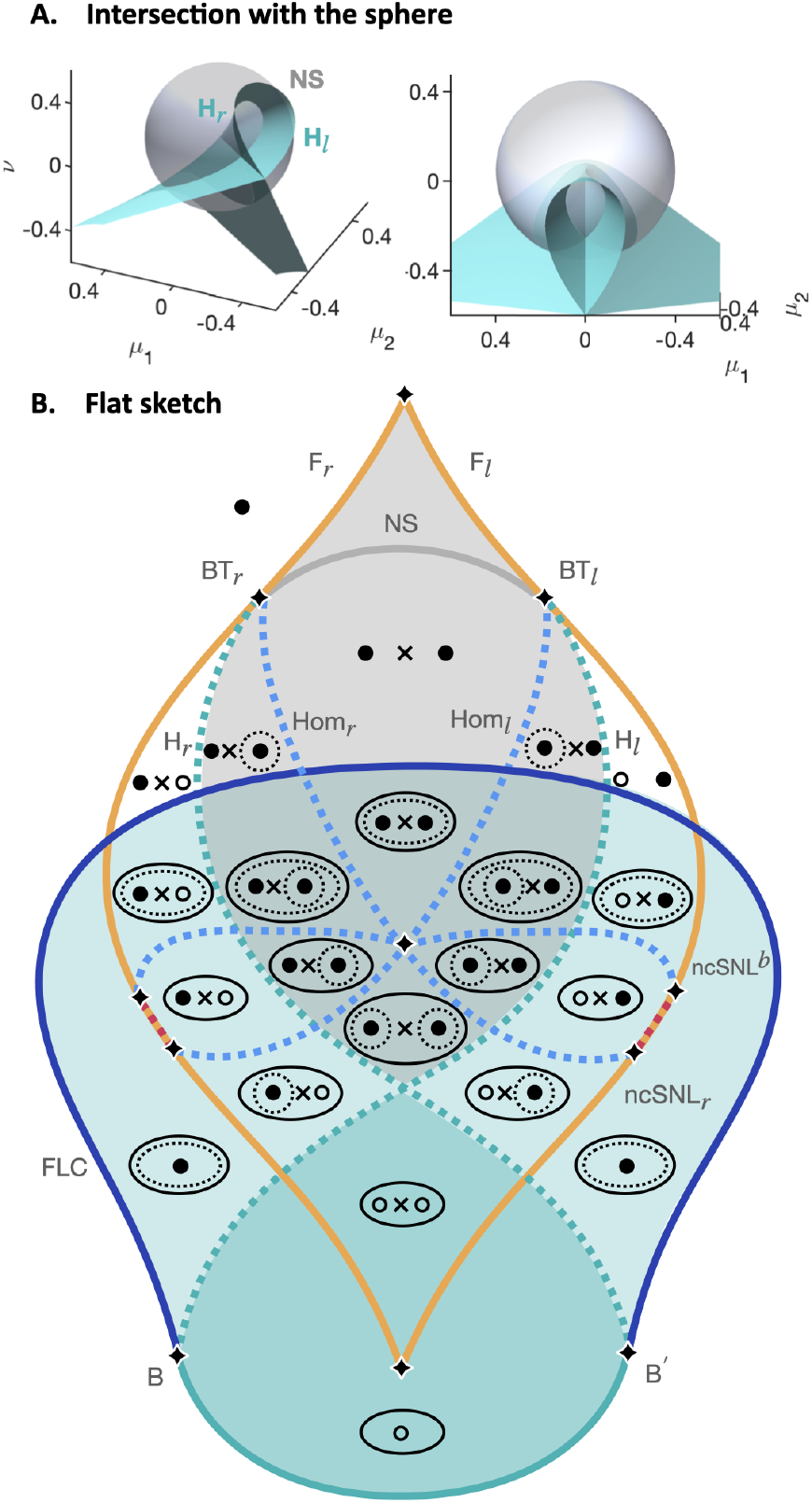
Unfolding the symmetric case on the sphere (b = 0). **A**. Intersection between the tr(*J*) = 0 manifold of Hopf and Neutral-Saddle and the sphere. **B**. Flat sketch of the bifurcation topology (adapted from [39]).

#### Intermediate cases in the literature

When fixing a positive *b*^*\**^ *>* 0, for *R > R*^*\**^ we encounter bifurcation diagrams with different topologies. If the topology changes, because of codim 3 bifurcations, we have transitioned to a different *case*. The conjecture is that, for big enough values of the radius, this sequence of intermediate cases leads to the topology of the symmetric one, that is, case (a). This sequence, and the main codim 3 bifurcations involved, have been hypothesized in [39]. Some of these cases have then been updated through numerical analysis in [41], which, starting from a small radius yielding the unfolding of the DBT-focus case (m), provided the subsequent five cases for increasing *R* (Fig. 4C, top). Continuing this exploration to reach the symmetric case would require increasing *R* towards infinity.

Equivalently, we can fix *R*^*\**^ *>* 0 and vary *b*. The bifurcation diagram of the unfolding of the DBT-focus is then found for a big enough *b*, and the symmetric case for *b* = 0. For intermediate values of *b*, the bifurcation diagram goes through the sequence of topologically different cases connecting the two extremes (Fig. 4C, bottom).

## 2 Results

To improve our understanding of the bifurcation structures arising from the unfolding of the DDBT bifurcation, we examine parameter space through spherical and planar sections.

### 2.1 Exploring with spherical surfaces

#### 2.1.1 A sequence of bifurcation diagrams linking the symmetric case to the DBT-focus one

As discussed above, numerically exploring the unfolding using spheres of different radii and fixed *b* led to the update of the conjectured cases closer to (m) [41]. However, the difficulty with further increasing *R* is that no lower bound is known for the value of the radius that guarantees that the topology of the symmetric case is reached. On the other hand, we know that for *b* = 1, the DBT-focus case can be found on spheres with *R <* 0.5 [56], while the symmetric case occurs as usual for *b* = 0. We can thus consider a fixed spherical surface centered at the origin in the (*µ*_1_, *µ*_2_, *v*) space and with *R* = 0.4 and explore the sequence of bifurcation diagram topologies on it for *b ∈* [0, 1], connecting the symmetric to the DBT-focus case. Topological sketches of the results are shown in (Fig. 7). We labeled cases in lowercase, while uppercase is used in [39, 41].

**Fig. 7.**
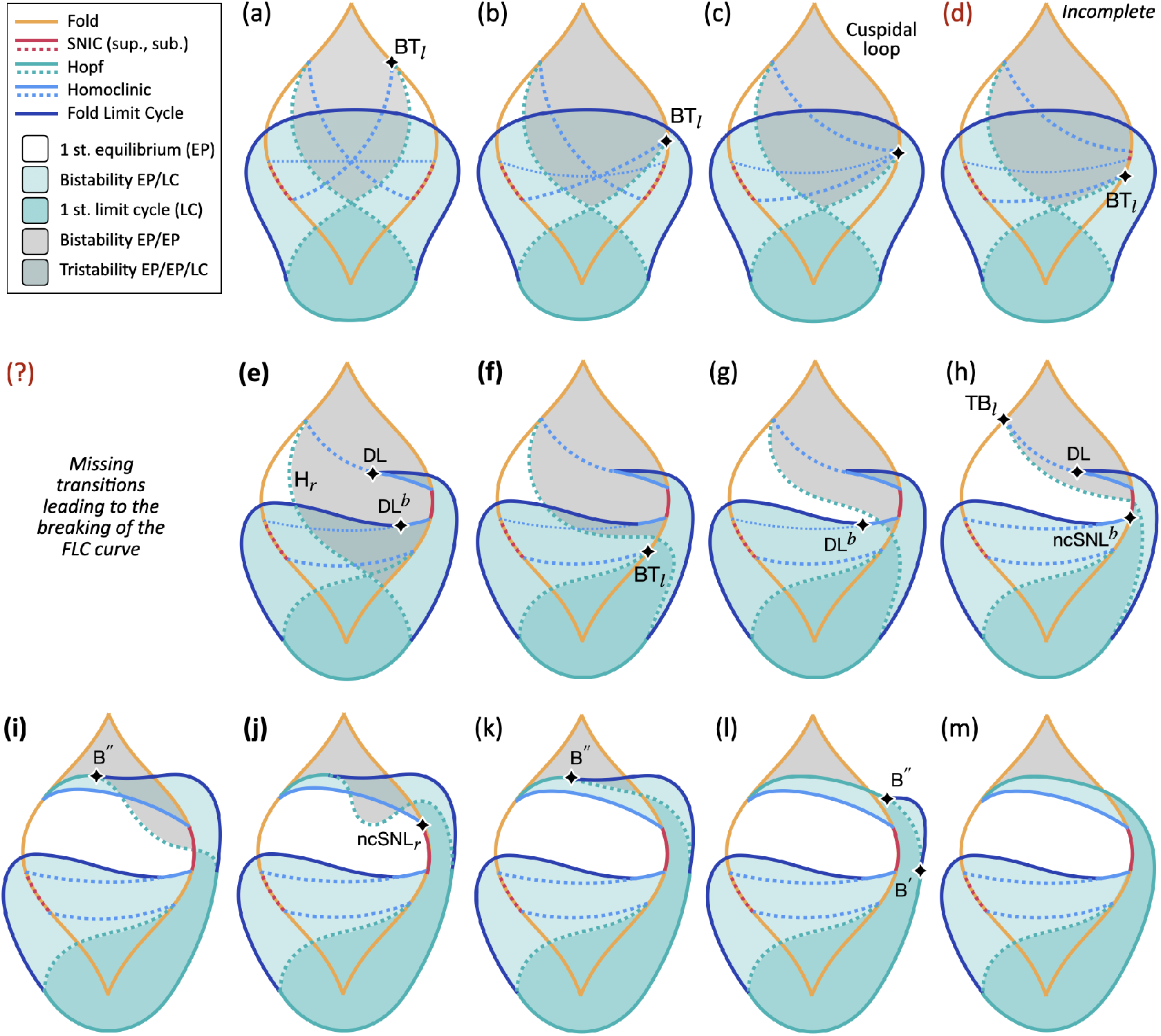
Varying *b* enables transitions between symmetric and DBT cases - numerical results and comparison with the conjecture. Topological sketches of the numerical results (the differential structure is not always preserved). When increasing *b*, the bifurcation diagram on the sphere loses the symmetry of the *b* = 0 case (a) and undergoes further modifications as bifurcation curves and points change their relative position with respect to one another [39, 41]. Topologically equivalent diagrams obtained for consecutive steps of *b* are grouped together in one *case*. In each case, we mark with a black diamond the codim 2 points involved in the transition to neighboring cases, and report their label. The colored background highlights how the regions allowing for different attractors deform during the transition. The *b* parameter does not alter the equilibria manifold (even though the stability of the solutions can change), nor the fold one, while the other curves are affected. A more detailed version of this figure is shown in Fig. 15. The Hom^*b*^ curve, when thinner, is not obtained numerically but based on [39] or consistent with neighboring cases. Cases in bold (e-f) differ from the conjecture [39], while (i-j) replace a case reported in a previous numerical study [41]. The bifurcations involved in the breaking of the FLC curve, conjectured in [39] and here confined between (d-e), remain to be verified.

We obtained the curves numerically, as detailed in Materials and Methods. We did not compute the Hom^*b*^ curve for cases (a-d). For these cases, we placed this curve according to the literature and consistently with the rest of the diagram, and we drew it thinner in Fig. 7.

Case (a) is the symmetric case studied in [39]. Subsequent cases occur when the diagram loses symmetry and curves change their relative position. Along these transitions, the values of *b* at which codim 1 curves and codim 2 points coincide, and separate two different cases, mark codim 3 events [39] (see the note on codimension at the end of this section).

Cases (b-c) are in agreement with the conjectured (B-C) in [39]: as *b* increases, the diagram loses symmetry with BT_*l*_ moving down. BT_*l*_ meets FLC (a codim 3 BT+FLC bifurcation) and then moves below it, giving the new topology in (b). When BT_*l*_ coincides with the right extremes of the other homoclinic curves (a codim 3 *cuspidal loop* [21]), we have the transition to (c). This latter event has been proven in [39].

Between cases (d-e), we observe the FLC curve breaking in two parts, creating two Degenerate-Loop (DL) points, one with Hom_*r*_ and the other with Hom^*b*^. In [39], these transitions (D-G in their paper) imply the appearance of a second FLC curve connecting the two homoclinic ones (through DL points) and containing a Cusp of Limit Cycles (CLC). As we only continue FLC curves from B points, we cannot confirm this mechanism. However, the behavior of the lower FLC branch after the breaking is different from what is entailed by the conjecture, as it does not show any CLC bifurcation on it, in accordance with [41] for later stages. From this numerical exploration, it appears, thus, that the secondary effects associated with the breaking of the FLC curve subside before case (e).

Once the FLC curve has split in two, in (e), the next transitions are caused by H_*r*_ going above BT_*l*_ in (f). These cases, which have not been proven before, differ from the conjecture due to the lack of the CLC. Case (e) relates to the conjectured (G), while (f) is implicit in [39].

The following topologies link back to the work of [41], in which cases equivalent to (g-l) lead to the final DBT case (m). The exceptions are (i-j), which are absent in [41]. In the latter work, the transition between and (k) occurs through a single different case (I’ in [41]). To corroborate our results, we performed the numerical continuation for these cases with other values of *b* and *R*, including those used in [41], and confirmed, for these choices, the occurrence of (i) and (j).

We now describe (g-l) in more detail. In (g), H_*r*_ moves above DL^*b*^, and above ncSNL^*b*^ in (h). Here, between (h) and (i), DL and BT_*r*_ coalesce. This brings to in which H_*r*_ and Hom_*r*_ are inverted, and DL disappeared, replaced by B^*′′*^. This transition occurs through a codim 3 DBT-cusp bifurcation.

In (j), H_*r*_ moves above ncSNL_*r*_, with a portion still below Hom_*r*_. In (k), H_*r*_ is fully above Hom_*r*_.

Lastly, B^*′′*^ moves rightward of *F*_*l*_ in (l), before the FLC curve disappears, through a codim 3 *non-transversal Bautin*, leading to (m).

To summarize, the original conjecture contained cases (A-M) [39], with (G-L) updated in [41] through numerical results and relabeled as (G’-H’-I’-K’-K”). Here, cases (B-D) are partially confirmed, except for Hom^*b*^, which was not computed; cases (E-F) are updated to the topologies shown in (e,f); and (I’) is replaced with two new cases (i-j). The breaking of the FLC curve is now confined to transitions between (d) and (e), but remains a key open problem.

#### 2.1.2 A note on codimension and geometrical bifurcations

For simplicity, in the above description, we have referred to a variety of events involving three simultaneous conditions being satisfied as codim 3 bifurcations, and we will maintain this notation in the remainder of the paper. However, it is worth noticing that there are differences among these codim 3 events due to whether the conditions apply to one or more objects in state space, and/or on how parameter space is explored.

Some of the codim 3 bifurcations mentioned above involve, in the most standard way, three conditions holding for a single object in state space. Examples are the DBT-focus and DBT-cusp bifurcations, where multiple degeneracies occur at a single equilibrium. These bifurcations are marked with filled diamonds in Fig. 7.

Other codim 3 events involve conditions on different objects in state space. These are referred to as codim “one-plus-two” bifurcations by Krauskopf & Osinga [41]. An example would be SNL + H, where the two bifurcations affect different equilibria.

Finally, there are cases in which the change in topology is due to the fact that the bifurcation manifold intersects the sphere non-transversally. These events have been recently formalized as *geometrical bifurcations* [3], because additional codimensions arise from the geometry of the bifurcation manifold and of the manifold used to explore the parameter space. An example is between stages (l-m) when the two codim 2 B points coalesce and disappear. This is a codim “two-plus-one” geometrical bifurcation, where now “two” refers to the codimension from a dynamical point of view (that of the Bautin bifurcation), and “one” from a geometrical point of view (spheres are a one-parameter family of surfaces used to probe the unfolding). Other examples will be discussed in Fig. 11.

We could thus consider both dynamical and geometrical codimensions, where the first can sum conditions on different objects in state space as in [41]. An example would be in (i-k), where the two intersections H+Hom coalesce and disappear (*dynamical* codim (1+1) + *geometrical* codim 1), giving another type of codim 3 event.

#### 2.1.3 The unfolding of the codim 4 DDBT and how it organizes codim 3 curves

We developed two new visualizations for a cohesive understanding of the several cases and transitions among them, summarizing known and new results.

We describe how codim 3 and codim 2 bifurcations vary with *b* by plotting, for each value of *b*, the values of the spherical coordinates *ϕ* of the codim 2 points found on the sphere (Fig. 8). Different *b* values are shown as concentric circles, with *b* = 0 at the origin, and *b* = 1 on the outer circle. This allows us to see, for each *b*, which codim 2 bifurcations appear, persist or disappear, as organized by codim 3 bifurcations. The missing bifurcations involved in the breaking of the FLC curve are reported from the original conjecture [39], but their position is confined between cases (d) and (e) consistently with the numerical results presented here and in [41]. More information can be found in the Supplementary Fig. S1.

**Fig. 8.**
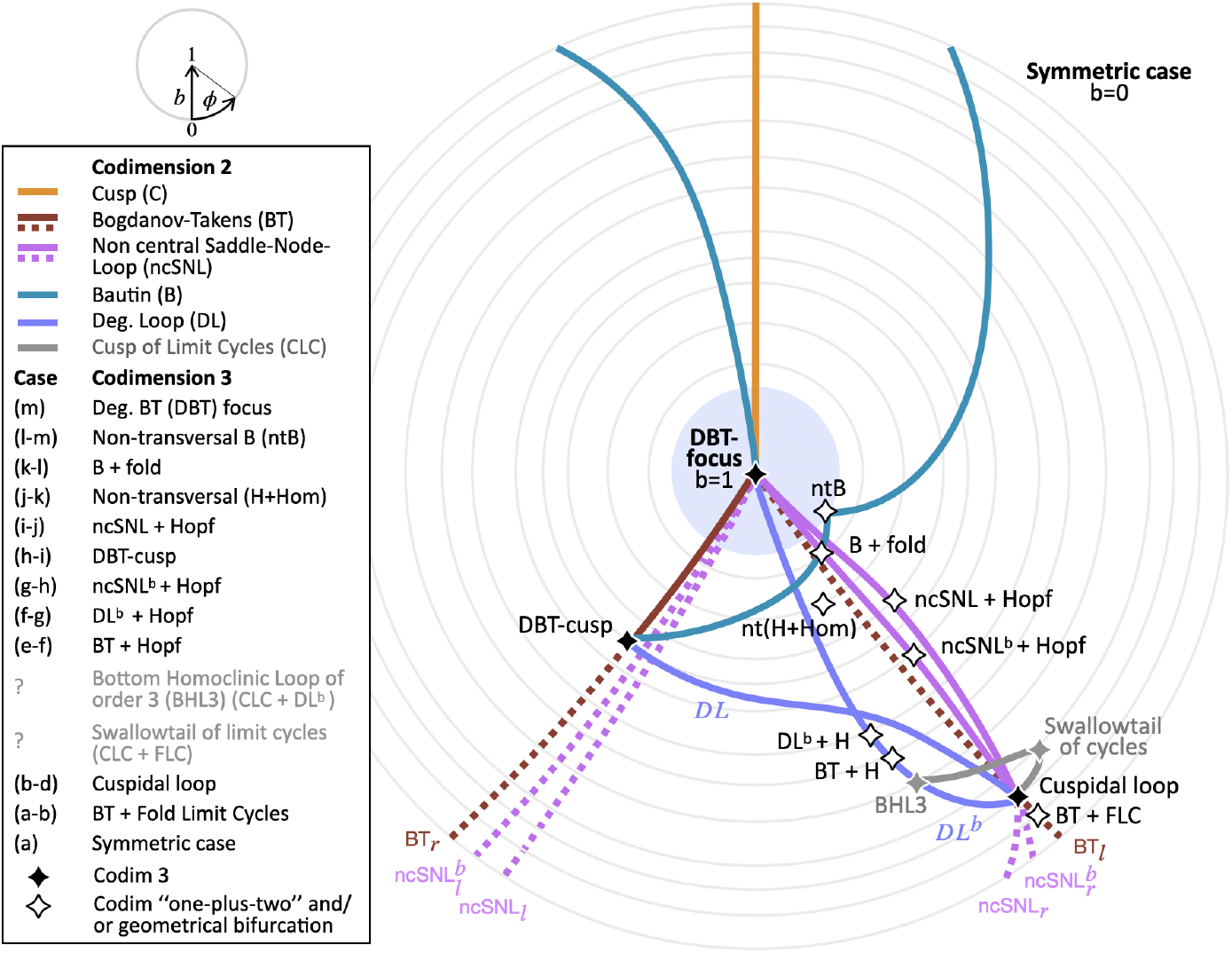
The codim 4 DDBT organizes lower-order bifurcation curves. Sketch based on the numerical results. Each circle corresponds to a value of *b*, from *b* = 1 at the center to *b* = 0 on the outer circle. For each *b*, we consider all codim 2 points on the sphere and plot their *ϕ* spherical coordinate on the circle. A light blue background marks the neighborhood of the DBT bifurcation where the cone structure is preserved. For the breaking of the FLC curve, we report, in gray, the two main events of the conjecture [39]. The Cusp bifurcation has spherical coordinate *θ* = 0, so that *ϕ* is undefined. We drew it here with *ϕ* = *π/*2.

The unfolding of the codim 4 DDBT, on the other hand, can be sketched by describing the three parameters (*µ*_1_, *µ*_2_, *v*) through the radius *R*, and drawing the (*b, R*) space (Fig. 9). Here, each point refers to a spherical surface, and we can observe how the topology of the bifurcation diagram on it varies through the parameter space, as dictated by the codim 3 bifurcation curves stemming from the DDBT bifurcation point.

**Fig. 9.**
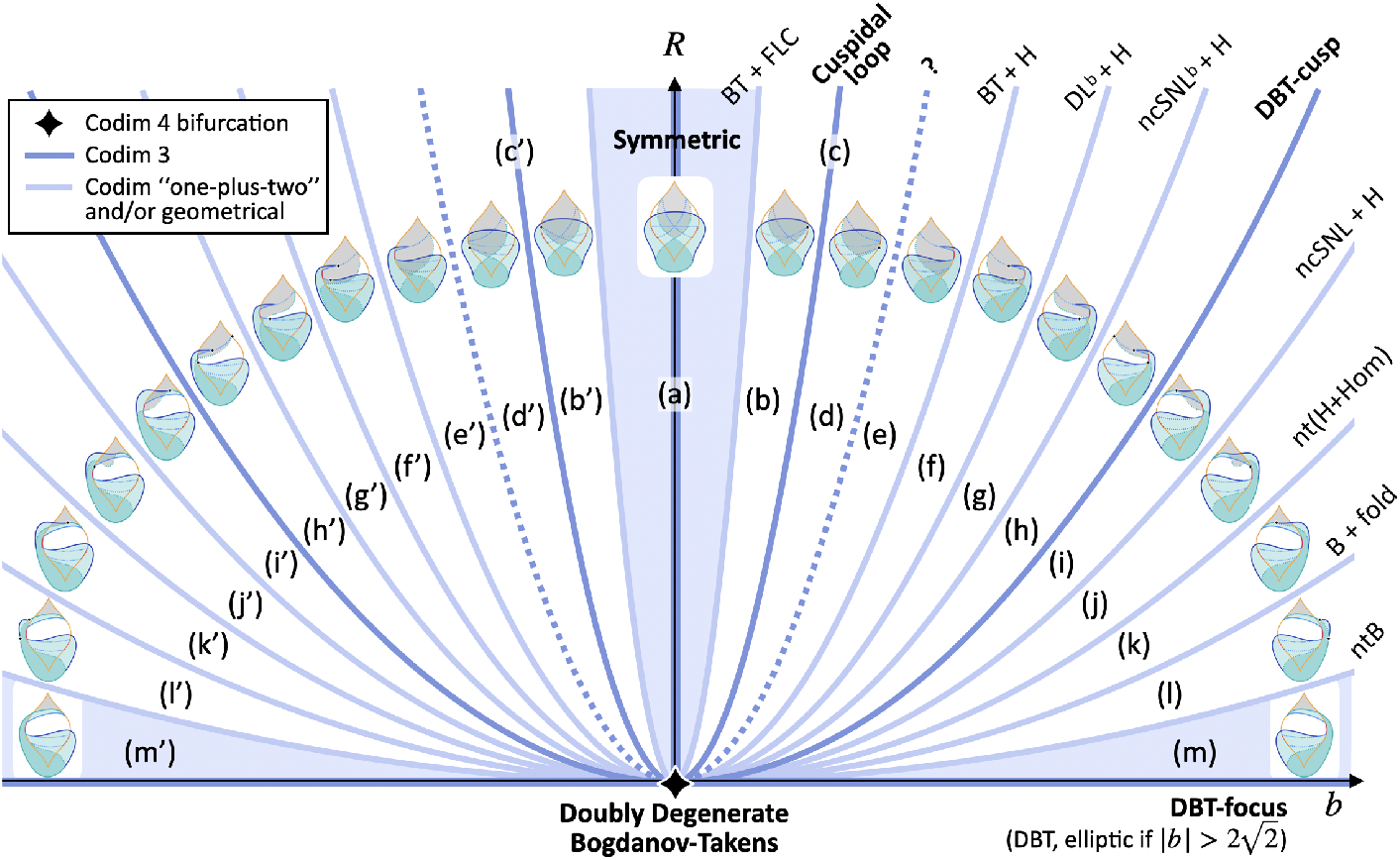
The unfolding of the DDBT bifurcation. The DDBT point is at the origin of the four-dimensional parameter space (*µ*_1_, *µ*_2_, *v, b*). We can visualize its unfolding by representing (*µ*_1_, *µ*_2_, *v*) collectively through the radius *R* of a sphere centered at the origin. Each point in the (*b, R*) plane then corresponds to the bifurcation diagram on a spherical surface of radius *R* for a given value of *b*. Codim 3 bifurcation curves stem from the DDBT bifurcation and partition the space into the different cases. The topologies are flipped for negative *b* (marked with a prime). A question mark reflects our incomplete knowledge about the transition from (d) to (e). The symmetric case has cone structure because the same topology persists for all values of *R* (light blue background, middle). The unfolding of the DBT-focus has cone structure in a neighborhood of the origin in (*µ*_1_, *µ*_2_, *v*) space (light blue background, bottom). The size of this neighborhood decreases with | *b*|. Taking a vertical cut in this map corresponds to exploring the unfolding through spheres of different radii and fixed *b*, while a horizontal one is the approach used here, with a fixed radius and varying *b*. Applying the method of concentric spheres to investigate this codim 4 unfolding would imply studying concentric half-circumferences in the (*b, R*) plane. This figure is a conceptual sketch.

### 2.2 New topologies found using planes

Because the bifurcation diagram of the intermediate cases lacks cone structure, the exact sequence of cases depends on the type of convex surfaces (spheres, parallelepipeds,…) chosen to explore the unfolding [41]. While closed surfaces, such as spheres, allow for a systematic exploration of the unfolding, in applications we can deal with other sections of parameter space. We thus explored whether planar cuts allowed for new topologies of bifurcation diagrams that could not be found on spheres.

Using concentric spheres, we can study the three-dimensional space (*µ*_1_, *µ*_2_, *v*) by changing one single parameter, the radius *R*. With planes, instead, we have more possibilities. The use of planes in the literature, so far, has been limited to planes with fixed *µ*_2_ [7, 26, 49], which implies the absence of cusp points, or for fixed *v* = 0 [49]. In this work, we focused on planes crossing the line of cusp bifurcations at a positive value of *ν* (in Fig. 2C this corresponds to the top cusp point) and intersecting ncSNL_*r*_, which plays an important role in neuronal models [34, 31]. We parametrized a plane through three points *P*_1_, *P*_2_, *P*_3_. The two coordinate vectors on the plane are 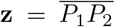 parallel to *−µ*_1_, and 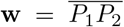 perpendicular to *µ*_1_. The parameter 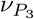, i.e. the *v* coordinate of point *P*_3_, determines the distance, in (*µ*_1_, *µ*_2_, *v*) space, between the cusp and the origin. However, because of how we defined the plane vectors, the cusp in the plane coordinates is always at (*z, w*) = (0, 1). This simplifies our numerical analysis and puts the focus on topology.

Interestingly, for different values of 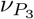 we find a range of *b* values for which the bifurcation diagram on the plane is topologically different from those we computed on spheres, as shown in Fig. 10. This is due to the H and/or Hom_*r*_ curves, stemming from BT_*r*_, bending back towards *F*_*r*_ because of how the plane intersects the codim 1 bifurcation manifolds. These new topologies cannot be explained by the missing cases on spheres (d). Rather, they involve different geometrical bifurcations.

**Fig. 10.**
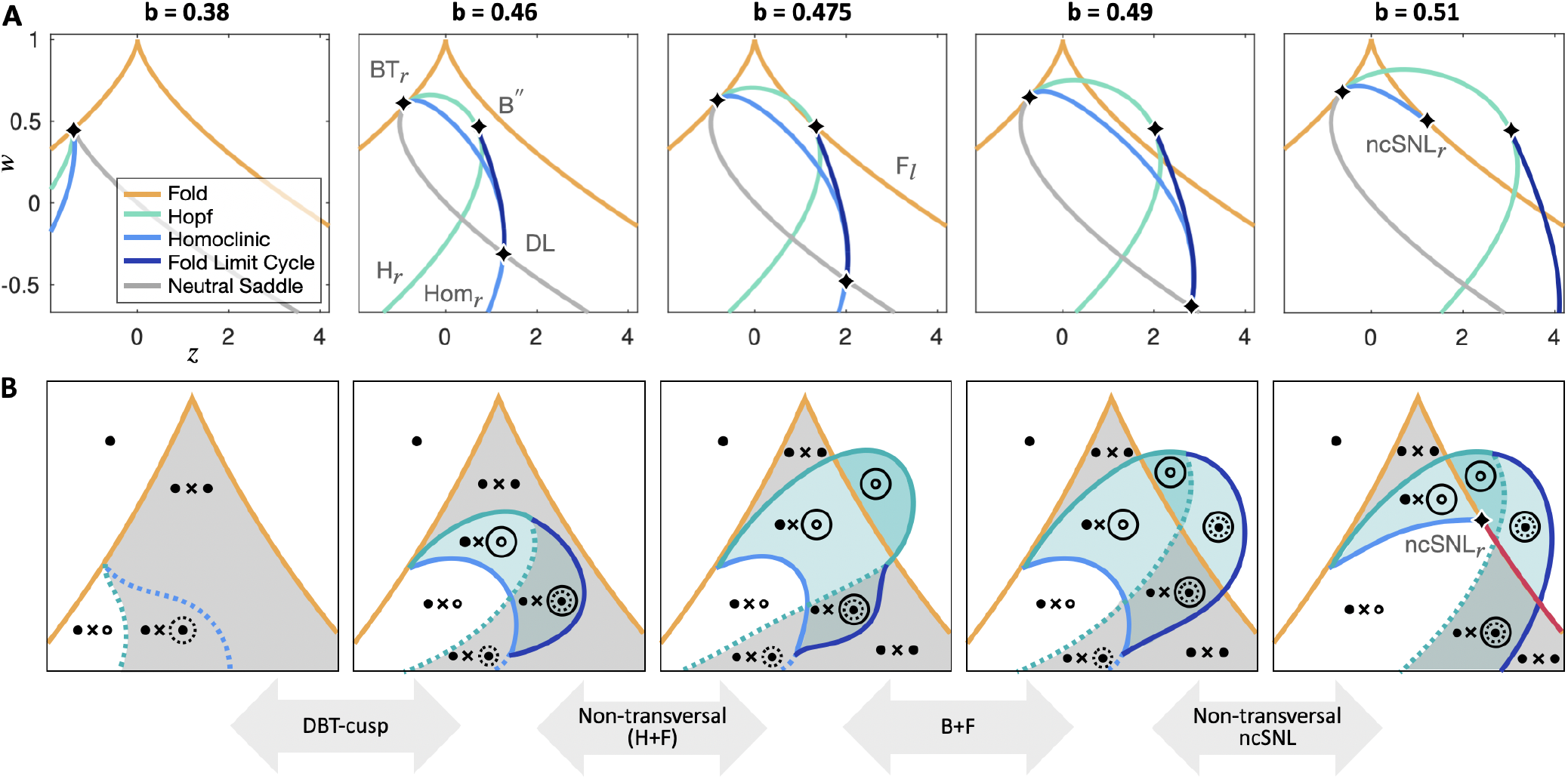
New topologies of bifurcation diagrams found on planes. **A**. We considered a plane in the (*µ*_1_, *µ*_2_, *v*) unfolding parameter space, parametrized by the two vectors **z** and **w**. When changing *b*, the bifurcation diagram includes new topologies that cannot be found on spheres. For these plots 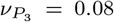. **B**. Sketches topologically equivalent to the corresponding bifurcation diagram of panel A. We include information about state space configuration and the codim 3 events involved in the transitions.

For small *b*, both H_*r*_ and Hom_*r*_ are subcritical. In the second panel, they become supercritical through the DBT-cusp bifurcation. The H_*r*_ intersects F_*l*_ in two points in the third panel (this change involves a geometrical bifurcation). The B” point then moves to the right of F_*l*_. Finally, Hom_*r*_ meets F_*l*_ forming ncSNL_*r*_ through another geometrical bifurcation. The SNIC curve below it can terminate in another ncSNL, which we did not detect.

The bending back of the curves is stronger for smaller values of 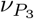 and less pronounced for bigger ones. For small 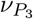, we can, however, always find a range of *b* producing these topologies.

Another consequence of this bending is that, on planes, we can cross the BT_*r*_ manifold twice. An example is shown in Fig. 11.

**Fig. 11.**
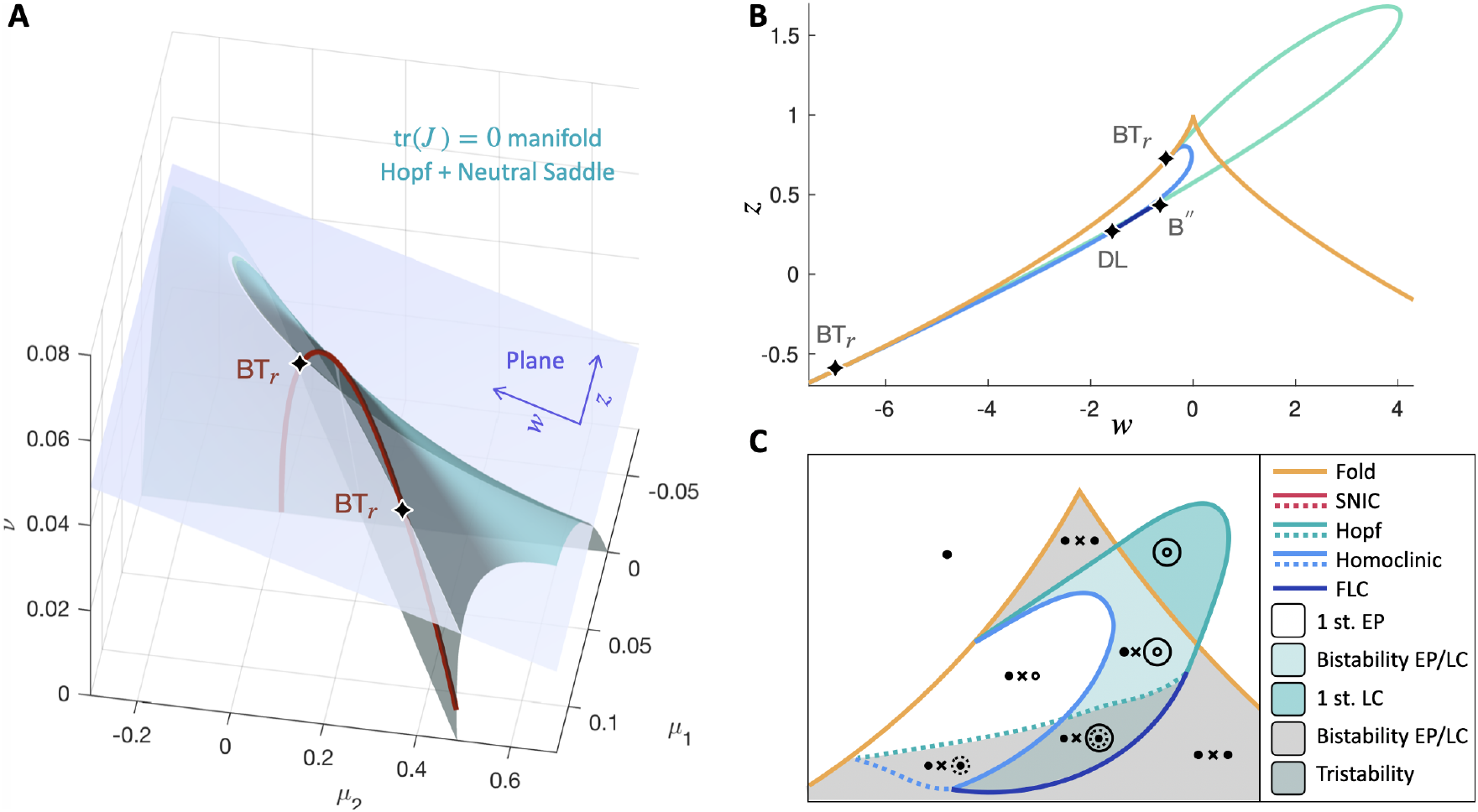
Planes can intersect the same codim 2 curve twice. **A**. A 3D view showing how some planes can intersect the same codim 2 curve, here BT_*r*_, more than once. This figure also provides an example to visualize geometrical bifurcations: if the plane is moved upwards by changing one parameter, the two BT_*r*_ on it first coalesce when the codim 2 curve and the plane meet non-transversally, and then disappear. This is a geometrical bifurcation because it depends on how the bifurcation manifold, here the codim 2 curve, meets with the one-parameter family of planes, that is, with the manifold used to explore the higher-dimensional bifurcation diagram. It has codim “two-plus-one” [3]. Parameters are set to: *b* = 0.42; 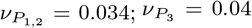. The non-transversal ncSNL brings to the appearance of two ncSNL_*r*_ for *b* = 0.46 (not shown). **B**. Bifurcation diagram on the plane for the situation of panel A. **C**. Topologically equivalent sketch.

### 2.3 Spheres and planes identify different transitions to excitability and bursting

Because transitions among bifurcation topologies found on planes can differ from transitions on spheres, due to geometrical bifurcations, behaviors relevant for a given system can appear/disappear in different ways in the two scenarios. As a proof of concept, we address transitions affecting (i) the SNIC curve, which is involved in neuronal excitability, and (ii) the possibility of supporting fold/hom bursting, a form of bursting common in many excitable cells.

When a system is at rest close to a supercritical SNIC, it can respond to small inputs with a large excursion in state space before setting back to rest. This is the dynamical mechanism for action potential generation for neurons with Type I excitability [54, 34], that is, neurons able to fire at arbitrarily low frequencies depending on the strength of an applied current. Fold/hom bursting instead is a type of fast-slow bursting. It requires a slow variable to steer the fast system back and forth along a path in its parameter space across fold and homoclinic bifurcations [34]. This burster exploits the bistability and feedback between fast and slow variables to alternate between resting and oscillatory behavior. Both these behaviors are organized by the ncSNL bifurcation [34, 26].

The sequences of topologies identified through spheres and planes show different transitions through which these behaviors can be impaired when parameters vary (Fig. 12). In bifurcation diagrams on spheres, ncSNL_*r*_ is present throughout the cases, but excitable behavior and fold/Hom bursting are only enabled after the cuspidal loop bifurcation has taken place. Both excitability and fold/Hom bursting first occur with the coexistence of a second stable equilibrium. From stage onwards, the second equilibrium becomes unstable. With planes, ncSNL_*r*_ only appears after the Hom_*r*_ curve meets *F*_*l*_. This enables excitability, with only one stable equilibrium. However, fold/Hom bursting is preserved before and after this transition.

**Fig. 12.**
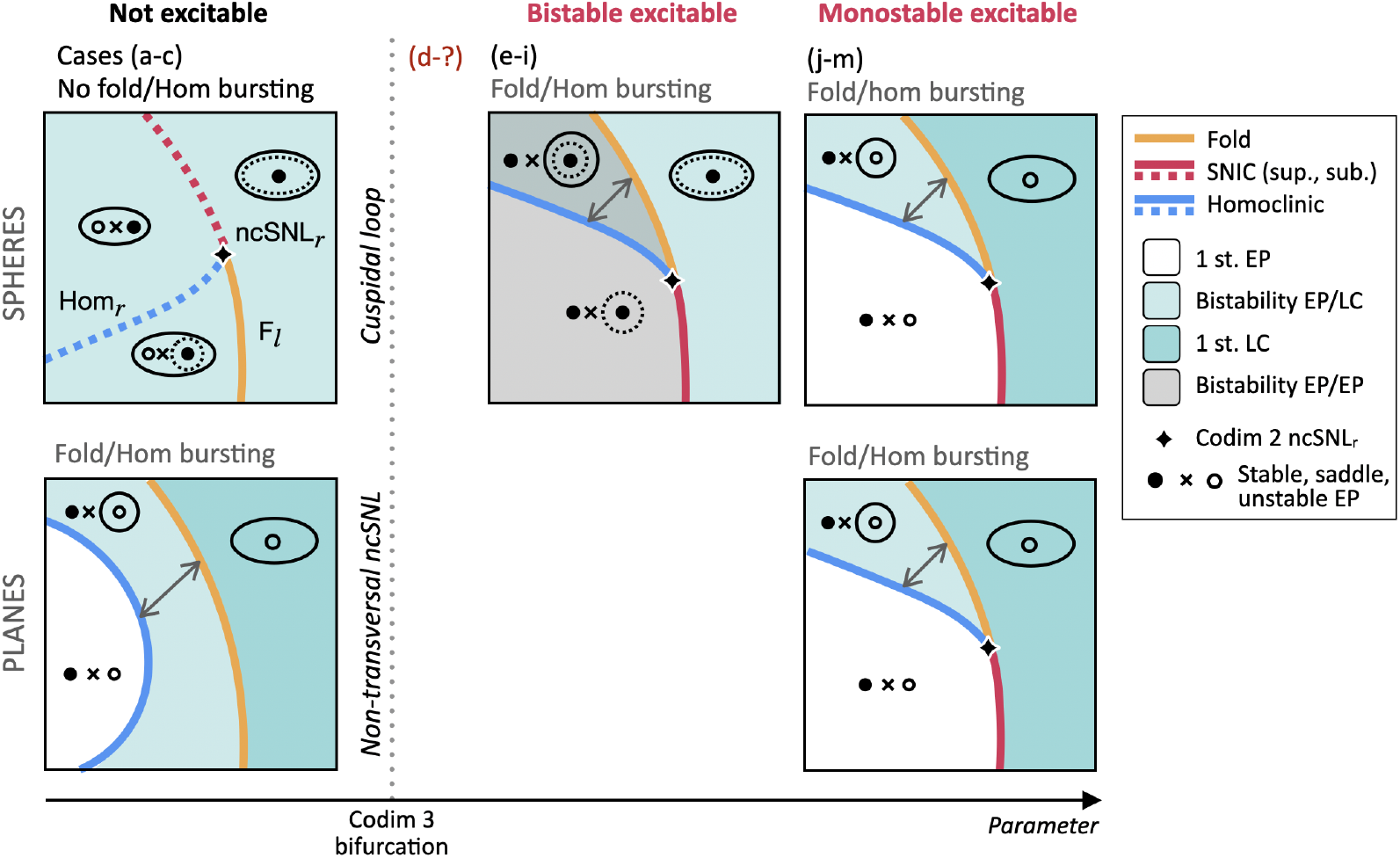
Spheres and planes identify different dynamical ways in which excitability and fold/hom bursting can be impaired. Proximity to a SNIC bifurcation enables excitable dynamics, that is, large transients in state space in response to small inputs when the system is at equilibrium [54, 34]. Fold/Hom bursting instead requires a slow variable steering the system back and forth across fold and homoclinic bifurcations (examples for such paths are shown as grey double arrows) [34]. These types of excitability and bursting, both relevant for excitable cells, are organized by the ncSNL bifurcation [34, 26], from which SNIC, fold and homoclinic curves unfold. The ncSNL bifurcation undergoes different codim 3 bifurcations when the unfolding is explored with spheres (top) or with planes (bottom), which alters how excitability and bursting are affected by parameter changes. With spheres, ncSNL_*r*_ persists through parameter variation, but both excitability and bursting are possible only after the cuspidal loop bifurcation, when the SNIC and Hom curve become supercritical. With planes instead, excitability is tied to the appearance of ncSNL_*r*_, through the non-transversal ncSNL bifurcation, while fold/Hom bursting is possible on both sides of this transition.

## 3 Discussion

We used spheres and planes to explore the unfolding of the DDBT bifurcations and identified some new bifurcation topologies. While using spheres is the standard approach, we show that planes add topologies that cannot be found and spheres, and discuss the potential biological relevance of the involved transitions in the study of the coexistence between neuronal Type I excitability and fold/Hom bursting.

### 3.1 Spheres

It has been conjectured that the unfolding of the codim 4 DDBT bifurcation connects the symmetric bifurcation diagram studied in [39] to that of the codim 3 DBT-focus type through a number of intermediate topologies. Some conjectured cases of the transition, those closer to DBT, have been modified after numerical exploration by Krauskopf and Osinga [41]. They probed the unfolding through spheres in a three-dimensional subspace, in which the DBT-focus lies at the origin, and increased the radius to detect the subsequent cases.

We here expand on that work and provide a numerical evaluation of other cases. To overcome the difficulty of dealing with increasingly bigger spheres and an unbounded range of radius values to explore, we exploit known properties of this unfolding [41] and consider spheres of fixed radius, while modifying the fourth parameter of the unfolding, *b*. The full range of transitions then lives in a bounded region of parameter space, that is *b ∈* [0, 1] for our choice of radius. We characterize all bifurcation diagrams except those for *b ∈* [0.29, 0.30], between cases (c) and (e) in Fig. 7, which are incomplete. Overall, we confirm most of the conjecture [39] and previous numerical results [39, 41] with differences in stages (e-f) and (i-j). However, the incomplete diagrams involve important conjectured transitions for the breaking of the FLC curve and remain the key open challenge.

The DBT case (m) appears in many neuronal models (see, for example, [40, 1]). With regards to the DDBT, the Morris-Lecar neuronal model [47], has, within its biologically relevant parameter range, a bifurcation diagram equivalent to case (h) (as shown in Fig. 6 of [29]). This provides another source of support, in addition to pseudo-plateau bursting [49], for the need to consider the full codim 4 unfolding to understand neuronal dynamics. Knowledge of the neighboring cases helps in anticipating which changes are to be expected when the model’s parameters are varied.

Beyond neuroscience, a bifurcation diagram topologically equivalent to the newly identified stage (i) has been proposed to encode the relationship between gene dosage and cell-division phenotype in a generic model for the regulation of the eukaryotic cell-cycle [12].

#### 3.1.1 Limitations and future directions

The main limitations of our work with spheres are that: we did not attempt the continuation of the Hom^*b*^ curve when the DL^*b*^ point disappeared (cases (a-d)); and we only continued FLC curves from Bautin points. These limitations constrained our understanding of the FLC breaking mechanism. In our numerical exploration, however, we do not see the secondary effects of the conjectured mechanism [39], such as the presence of CLC bifurcations, from case (e). This extends the observations of [41] to later cases. More details about the conjectured FLC breaking and our partial numerical results for it can be found in Fig. S1 in the Supplementary Materials.

It is interesting to note that a CLC appears in the bifurcation diagram of a well-known mesoscopic model for neural activity, the Jansen-Rit model [36] (see Fig. 4 in [59]), for which the DBT bifurcation has been identified. The CLC enables a distinctive feature of this model, that is, the coexistence of limit cycles with different amplitudes. A clarification of the transitions from the cuspidal loop to the FLC breaking will improve our understanding of such models.

Our results constrain these events to a precise range of *b* values, but the study of this transition will require more finely tailored approaches, which are beyond the scope of this paper. As discussed in [41], most of the codim 3 bifurcations involved between (d) and (e) appear in a closely related system [13], studied in terms of perturbations from a Hamiltonian system [39]. Future work to address (d-e) will require a combination of numerical and analytical tools. Numerically, for example, while Matcont cannot compute cuspidal loops, strategies exist to avoid this limitation (see, for example, [53]). Analytically, an iterative procedure for a perturbative solution to approximate cuspidal loops up to any wished order has been recently proposed and tested on two systems related to the normal form of degenerate BT bifurcations [53, 13, 24]. These can serve as starting points to tackle the missing transitions.

### 3.2 Planes

The use of planes to explore the unfolding leads to new topologies due to the inward bending of the Hopf and related bifurcation curves. We illustrated some implications of the new topologies for transitions to topologies able to support two biologically relevant behaviors: Type I excitability, a type of neuronal excitability shared by most cortical pyramidal neurons [35], which requires a supercritical SNIC [54]; and fold/Hom bursting, a type of bursting common in neuronal models and exhibited by a variety of excitable cells [34]. We showed that, while on topologies across spheres excitability and bursting were only possible for the same radius values, the transition across planes allowed for bursting to be possible even when excitability was impaired (Fig. 10).

The extreme bending of the Hopf curve involves bifurcation diagrams with two BT_*r*_ points (Fig. 11). In this scenario, the SNIC segment on F_*l*_ can be preserved, when Hom_*r*_ reaches F_*l*_ (see also [49]). The SNIC segment is instead lost, and thus the possibility of excitable behaviors close to it, if Hom_*r*_ bends back earlier, as in the figure. Diagrams of this type, with two BT points on the same fold branch, have been observed in the neural dynamics literature, for example, in the Wilson-Cowan model for the mesoscopic activity of a neural mass [63,9].

#### 3.2.1 Limitations and future directions

The use of planes is exploratory rather than systematic, and other topologies may be possible that we did not identify. We focused on the upper part of the unfolding and the role of ncSNL for excitable behaviors [35, 31, 30], but it has been shown that the lower portion of the unfolding supports a structured sequence of transitions in neural excitability Type as the system moves beyond codim 2 points [40, 1]. Future work will address whether new topologies in the family of planes intersecting the lower Cusp (negative 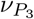) affect these transitions.

### 3.3 Conclusions

In neural dynamics, a wider acknowledgment of the presence of the codim 3 DBT bifurcation in models is helping to unify different excitable and bursting behaviors [49, 40, 56, 48]. The new topologies and transitions provided here will allow further investigation of the role of the fourth parameter of the DDBT in systems, also beyond neuroscience, for which DBT singularities have already been identified. Future work will systematically investigate the implications for excitable behaviors and fast-slow bursting.

## 4 Materials and Methods

We used Matcont version 7p4 and 7p6 [17] command line for numerical continuation.

### 4.0.1 Bifurcation diagrams on spheres

We expressed the parameters (*µ*_1_, *µ*_2_, *v*) in spherical coordinates (*R, θ, ϕ*) in Eq.1. We fixed *R* = 0.4 based on previous work [56].

The fold manifold does not depend on *b*, so its intersection with the sphere remains as in Fig. 2C (analytical results). The Hopf manifold, instead, is symmetric for *b* = 0 and loses symmetry while *b* increases (Fig. 14, analytical results).

For the numerical exploration, we varied *b ∈* [0, 1] with a step of 0.02. We initialized the system in the upper cusp, which can be shown analytically to occur for (*x, y*) = (0, 0) and (*θ, ϕ*) = (0, 0). We continued the fold curve (LP in Matcont), on which the two BT points were identified. From the BT points, we continued the Hopf (forward and backward) and homoclinic curves. If the Hopf curve had Bautin points (GH in Matcont), we continued FLC (LPC in Matcont) from there. We also followed homoclinic curves from the end point of the FLC ones to reconstruct Hom^*b*^ for cases from (e) onward. This was the most delicate continuation, which sometimes failed. We reconstructed this full curve at least once within a given case, except in (e-g), where the subcritical branch is placed according to subsequent cases, while the supercritical branch is nu-merical. We did not attempt the reconstruction of this curve before the breaking of the FLC curve. In those sketches, Hom^*b*^ is based on the literature, with that in (a) verified numerically in [39].

Our algorithm computes some curves several times (for example, forward and backward Hopf branch from the two BT points coincides, as do the Bautin points on them from which we continued FLC curves, and so on). A minimal implementation would be: from the cusp point, continue the folds; continue homoclinic curves from both BT points, but Hopf from only one of them (backward and forward); continue FLC curves from all Bautin points on the Hopf curve; continue the Hom^*b*^ from the end point of the lower branch of the FLC curve. However, computing the same curves with different starting points gave robustness to our results, in particular for delicate continuations such as the Hom^*b*^ one, so we maintained the redundancies.

We performed exploration at a higher *b* resolution to resolve transitions among cases when not clear. For the transition between (h) and (k) we used a step of 0.005 for *b ∈* [0.7, 0.75]. At this resolution, the transition between (h) and (k) was achieved through two intermediate cases, (i) and (j) (Fig. 13A), both different from the single transition case labeled (I’) in [41]. In the latter work, the unfolding was explored by fixing *b* = 1 and varying *R*. We thus fixed *b* = 1 and explored *R ∈* [0.75, 0.85], with step 0.01, which yielded again to (i) and (j) (Fig. 13B).

**Fig. 13.**
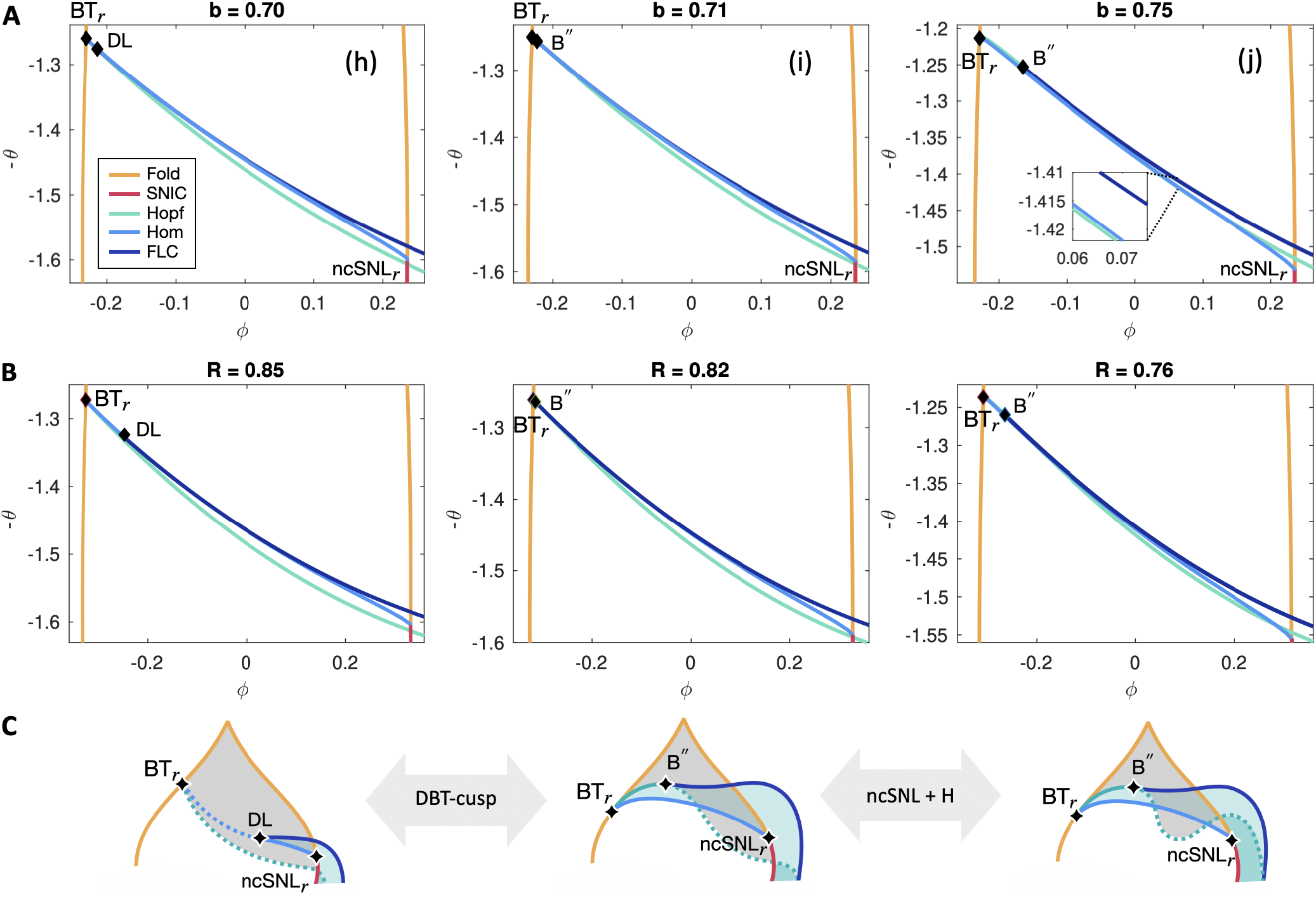
New cases in the transition from (h) to (j). Numerical continuation results. **A**. Fixed *R* = 0.4. In (h), the Hopf curve stemming from BT_*r*_ stays below the Hom_*r*_ curve. In (i), the first portion of H is above Hom_*r*_ and forms a Bautin point with FLC; it then passes below Hom_*r*_. In (j), H behaves as in (i), except going above Hom_*r*_ again before the latter disappears in ncSNL_*r*_. **B**. The same cases can be found by fixing *b* = 1 and varying *R*. **C**. Sketches of the topologies.

**Fig. 14.**
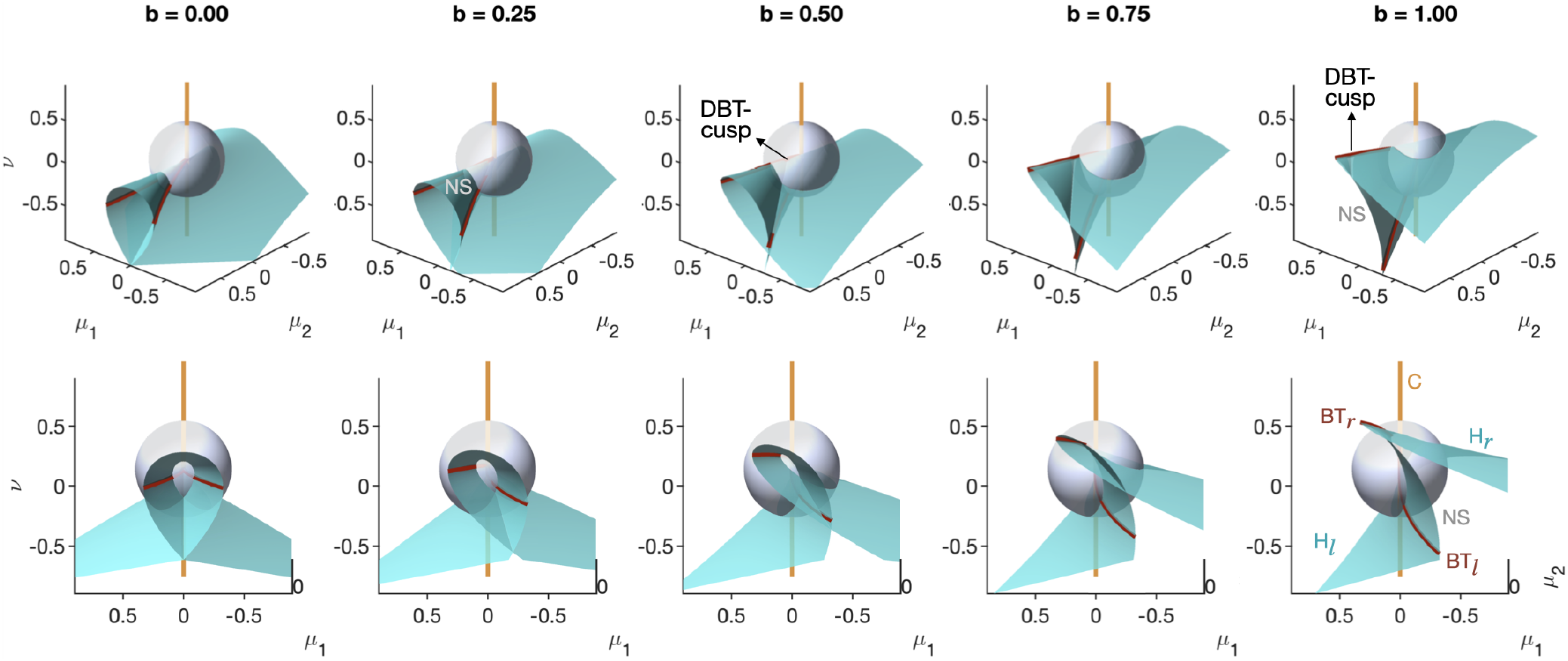
The Hopf manifold loses symmetry with varying *b*. Analytical results for the manifold corresponding to tr(*J*_*ep*_) = 0, which includes Hopf and Neutral-Saddle, which loses symmetry with increasing *b*. The NS portion, for small *b*, joins the two BT curves (dark red) on the top. When the codim 3 DBT-cusp point is outside the sphere, NS on the spheres joins the two BT points in the middle branch of the manifold. Two views, lateral (top row) and frontal (bottom).

For the transition between (d) and (e), where the FLC curve breaks in two, we used a step of 0.005 with *b ∈* [0.25, 0.32]. We also decreased step sizes and increased the sensitivity of the numerical continuation for the FLC curve. With this increased sensitivity, however, the numerically computed curve is already broken at *b* = 0.25, with the two branches overlapping. For *b* = 0.29, the two endpoints nearly coincide (arrow in Fig. S1) and separate for *b* = 0.3. However, one of the two endpoints does not coincide with the DL^*b*^ point, calling for improved continuation. In general, the FLC curve stemming from B ended before reaching the Neutral-Saddle curve. This may explain our difficulties in continuing the subcritical branch of Hom^*b*^ from there.

Examples of *b* values relative to each case are: (a) *b* = 0, (b) *b* = 0.27, (c) 0.265 *< b <* 0.27, (d) *b* = 0.28, (e) *b* = 0.32, (f) *b* = 0.34, (g) *b* = 0.48, (h) *b* = 0.64, (i) *b* = 0.71, (j) *b* = 0.75, (k) *b* = 0.8, (l) *b* = 0.9, (m) *b* = 1.

Fig. 15 provides a more detailed version of the cases, with a sketch of state space for each region.

**Fig. 15.**
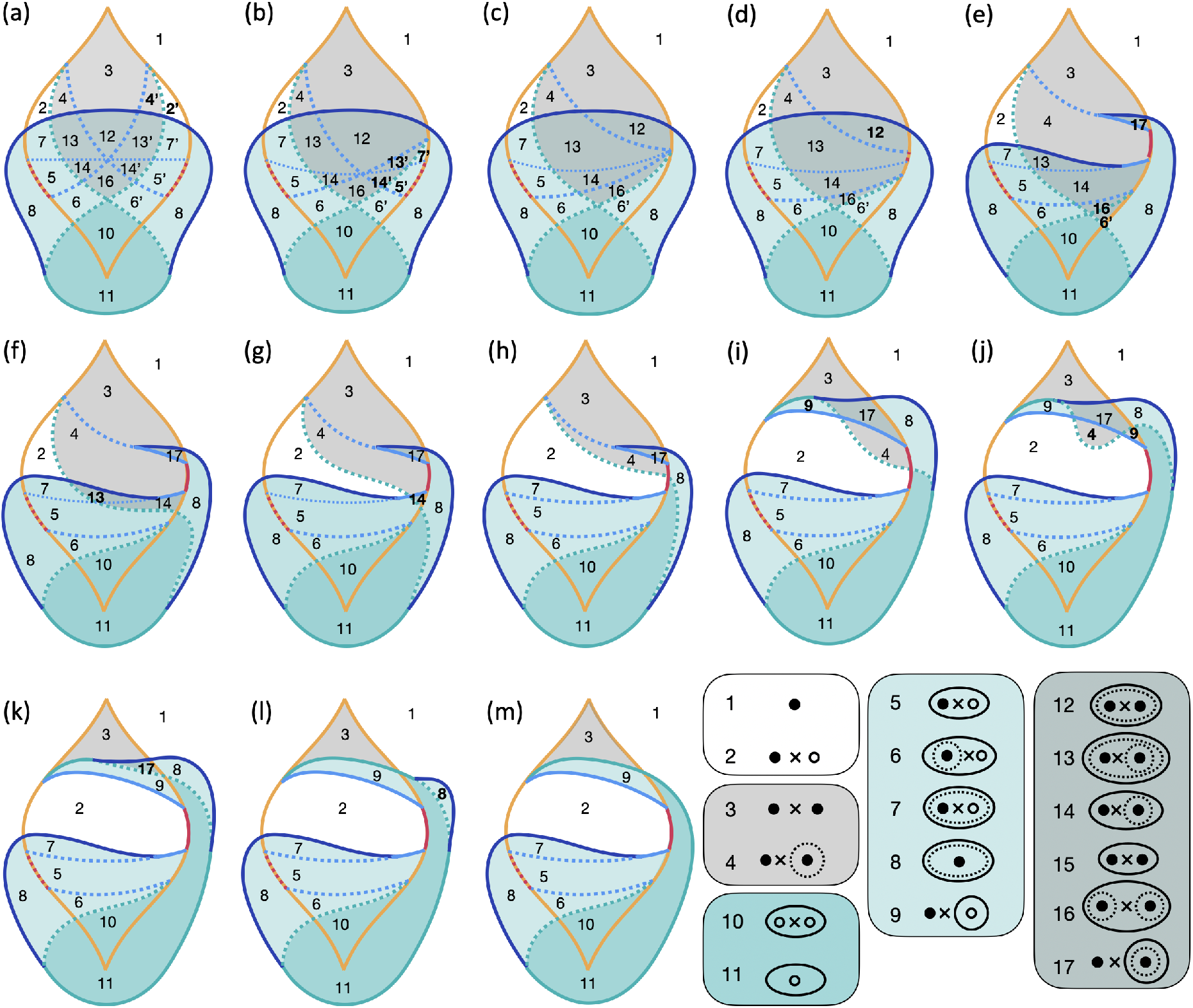
Transitions with information about state space configuration. This is a more detailed version of Fig. 7, where numbers in each region refer to a specific topology of stable/unstable equilibria (filled/empty dots), saddles (x) and stable/unstable limit cycles (full/dashed ellipses). Bold numbers mark regions appearing or disappearing in the neighboring cases. The background highlights the type of attractors available in each region: one equilibrium (white), two equilibria (grey), one limit cycle (teal green), one equilibrium and one limit cycle (light teal green), or two equilibria and one limit cycle (grey-green).

### 4.0.2 Bifurcation diagrams on planes

We considered the parametric equations of a plane through three points *P*_1_, *P*_2_, *P*_3_. We chose the points so that the plane would intersect the upper C point. We anchored *P*_1_, *P*_2_ such that 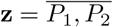 would be parallel to the *−µ*_1_ axis and have *µ*_2_ = 0.2. *P*_3_, instead, lies on the *v* axis at 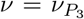. All figures use 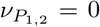 unless stated differently in the caption. This gives 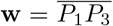 perpendicular to **z** and passing through C. We used (*z, w*) as bifurcation parameters on the plane and considered different 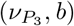 values to study how they affect the bifurcation diagram (Fig. 16).

**Fig. 16.**
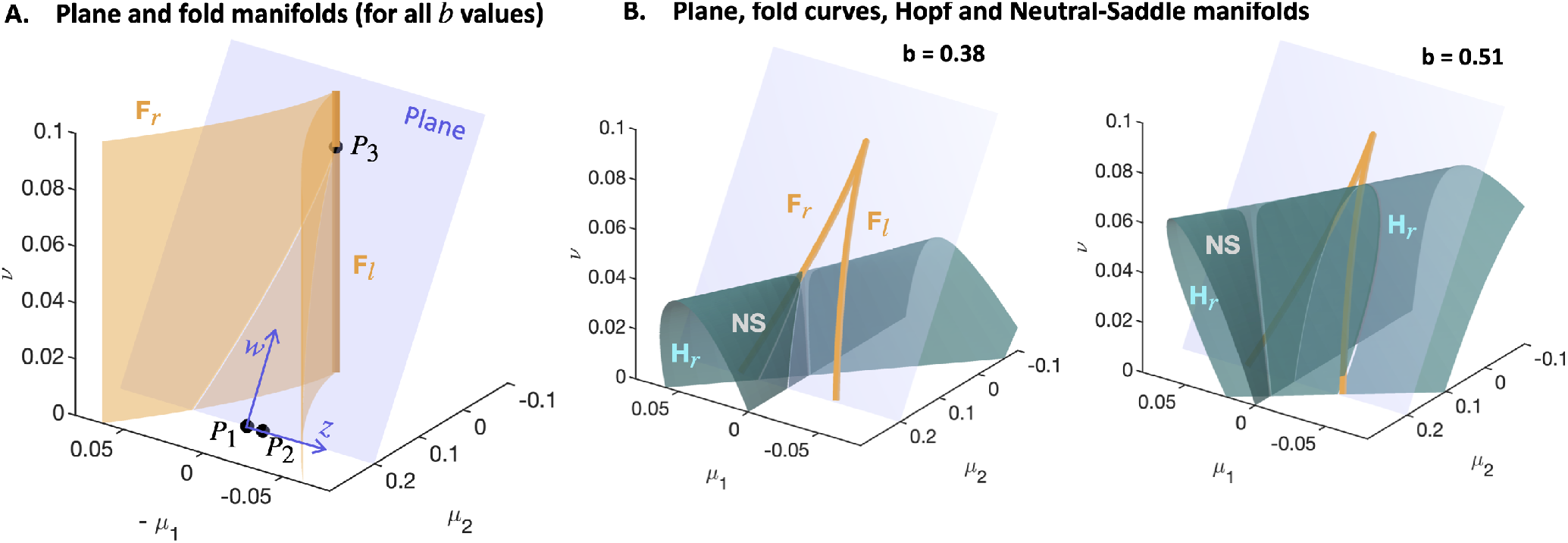
Construction of the plane and intersections with local codim 1 bifurcation manifolds. **A**. Plane through three points and intersection with the fold manifold. By construction, this intersection is the same for different values of *b* and 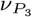 . **B**. Intersection with the manifold where the equilibria satisfy tr(*J*_*ep*_) = 0. This manifold represents H bifurcation and NS condition. We show the intersection with the plane for some of the values used in Fig. 10.

With our choices 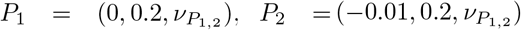 and 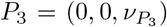, the plane’s parameterization reads:

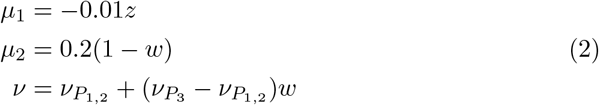

which we replaced in Eq.1. We initialized the system in the cusp, which can be found analytically to have (*x, y*) = (0, 0) and (*z, w*) = (0, 1). From there, we continued the fold curves, identified BT points and used them to continue Hopf and homoclinic. If the Hopf curve had a Bautin point, we continued the FLC curve from there. We did not compute other curves, such as Hom^*b*^, and we did not identify ncSNL^*b*^.

## Acknowledgements

I thank Viktor Jirsa, Jonathan Rubin and Tommaso De Lorenzo for stimulating discussions about this unfolding, and Charlotte Maschke and Gabriele Casagrande for providing valuable feedback on the manuscript. I am also grateful to the anonymous reviewers for carefully reading this work and suggesting many improvements and clarifications.

## Author contribution

MS is the sole author of this manuscript.

## Footnotes

1 ^1^ There are different versions in the literature with regards to the signs in Eq. 1, which can be obtained from one another by reversing the direction of *t, x*, or the parameter’s axes.

