## Supplemental Figure 1 for "New topologies in the unfolding of the Doubly Degenerate Bogdanov-Takens bifurcation"

**Noname manuscript No.**
(will be inserted by the editor)

### New topologies in the unfolding of the Doubly Degenerate Bogdanov-Takens bifurcation - Supplementary Material

**Marisa Saggio**

Received: date / Accepted: date

Marisa Saggio
Aix Marseille Univ, INSERM, INS, Inst Neurosci Syst, Marseille, France
Orcid: 0000-0002-3846-6100


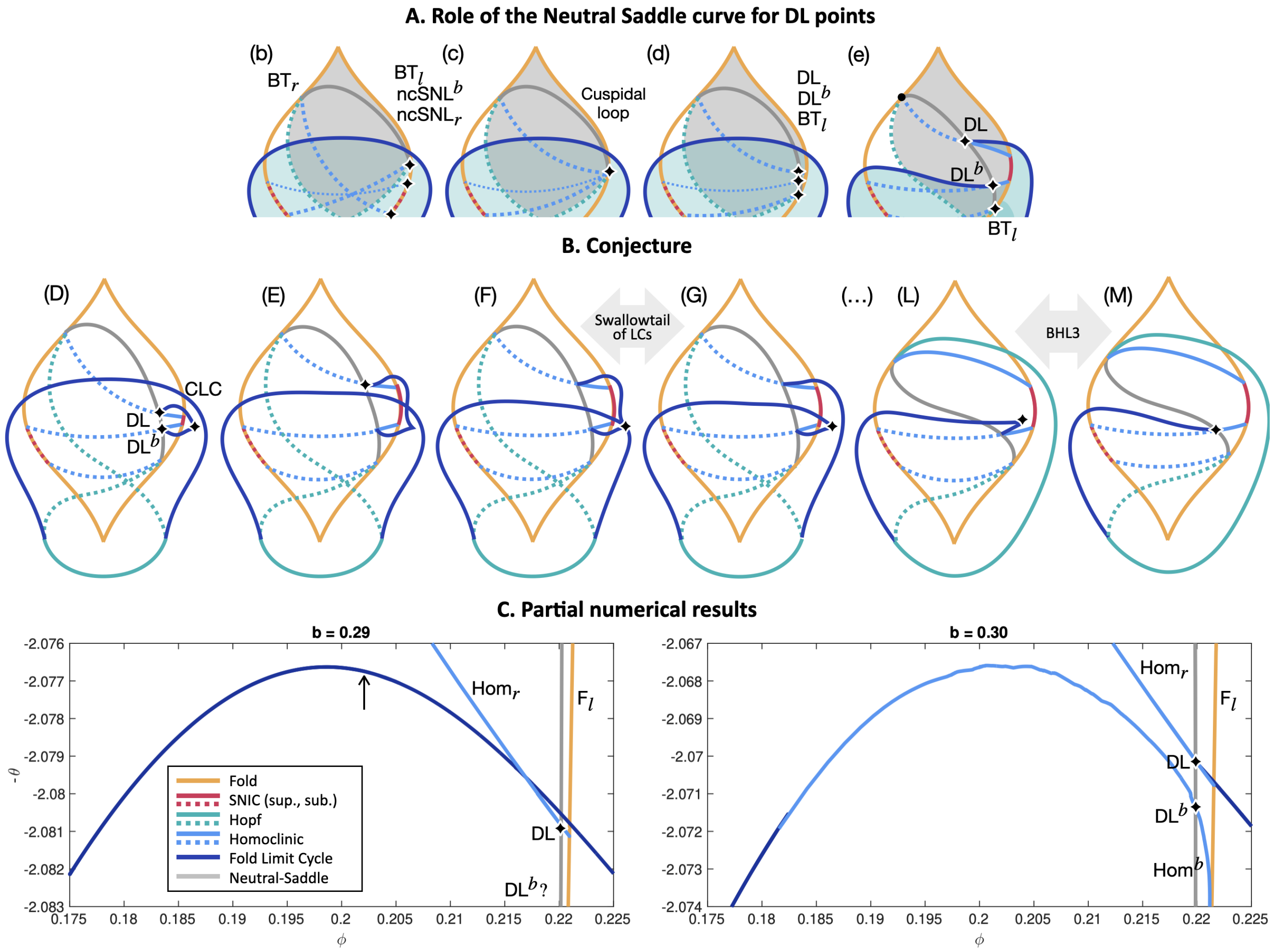


**Fig. 1 The breaking of the FLC curve. A.** Topological sketches. At a DL point, the homoclinic curve changes from being subcritical to supercritical or vice versa. This occurs when the saddle equilibrium becomes a *neutral saddle*, that is, when its *saddle quantity* (the sum of its two real eigenvalues) is zero [3]. DL bifurcations, as found in cases (e-l), thus occur at the intersection between the NS curve and the Hom curves. These intersections only exist when $BT_l$ moves below the ncSNL points, after (c). **B.** Topological sketches of the conjecture adapted from [2]. The author propose that a FLC curve joins the two DL points contains a CLC, as it follows from the unfolding of the codim 3 cuspidal loop [1]. This gives a limit cycle of multiplicity four when it intersects the old FLC curve (F), unfolded by a swallow tail of limit cycles, allowing the breaking of the FLC curve, and leaves a CLC on the lower branch, which persists until case (L). The CLC disappears through a bottom homoclinic loop of order 3, leading to (M). **C.** Numerical results in the parameter range where we observe the breaking of the FLC curve. $NcSNL^b$ from $DL^b$ was not computed in the left panel. An arrow points to where the FLC branches partially overlap in the numerical results. Intermediate stages are missing to connect the two bifurcation diagrams. While we cannot confirm or not the presence of the new FLC curve and its transitions, two differences from the conjecture appear: ncSNL seems to be below FLC when the curve breaks; no CLC is detected on the broken FLC branches. A clarification of this transitions will require more finely tailored approaches.
